# Gender impacts the relationship between mood disorder symptoms and effortful avoidance performance

**DOI:** 10.1101/2022.06.21.497075

**Authors:** Brandon J. Forys, Ryan J. Tomm, Dayana Stamboliyska, Alex R. Terpstra, Luke Clark, Trisha Chakrabarty, Stan B. Floresco, Rebecca M. Todd

## Abstract

We must often decide how much effort to exert or withhold to avoid undesirable outcomes or obtain rewards. In depression and anxiety, levels of avoidance can be excessive and reward-seeking may be reduced. Yet outstanding questions remain about the links between motivated action/inhibition and anxiety and depression levels, and whether they differ between men and women. Here we examined the relationship between anxiety and depression scores, and performance on effortful active and inhibitory avoidance (Study 1) and reward seeking (Study 2) in humans. Undergraduates and paid online workers (*N_Avoid_* = 545, *N_Reward_* = 310; *N_Female_* = 368, *N_Male_* = 450, *M_Age_* = 22.58, *Range_Age_* = 17-62) were assessed on the Beck Depression Inventory II (BDI) and the Beck Anxiety Inventory (BAI) and performed an instructed online avoidance or reward-seeking task. Participants had to make multiple presses on active trials and withhold presses on inhibitory trials to avoid an unpleasant sound (Study 1) or obtain points towards a monetary reward (Study 2). Overall, men deployed more effort than women in both avoidance and reward-seeking, and anxiety scores were negatively associated with active reward-seeking performance based on sensitivity scores. Gender interacted with anxiety scores and inhibitory avoidance performance, such that women with higher anxiety showed worse avoidance performance. Our results illuminate effects of gender in the relationship between anxiety and depression levels and the motivation to actively and effortfully respond to obtain positive and avoid negative outcomes.

**Significance statement:** We must often take or withhold effortful action to avoid unpleasant outcomes or obtain rewards. Depression and anxiety can impact these behaviours’ effectiveness, but the roles of avoidance in depression and reward-seeking in anxiety are not fully understood. Gender differences in avoidance and reward-seeking have also not been examined. We present a task in which community participants with a range of anxiety and depression levels made or withheld button presses to avoid hearing an unpleasant sound or obtain a reward. Men deployed more effort than women in avoidance, and women with higher anxiety scores had lower avoidance performance than men. We illuminate gender differences in how depressive and anxiety scores impact our ability to avoid threats and obtain rewards.

## Introduction

### Avoidance and reward-seeking behaviours

Living organisms are motivated to avoid potential threats or to acquire rewards respectively. Often achieving these goals requires action, but it can also require refraining from action. For example, we may take action to remove a threat’s potential harm through active avoidance, or we may decide that withholding action is the best way to let the threat pass by, as in inhibitory avoidance (Krypotos, Effting, Kindt, and Beckers, 2015; LeDoux, Moscarello, Sears, and Campese, 2017). Alternatively, in a situation that offers the possibility of reward, we may take action to approach the reward through active reward-seeking or, instead, inhibit pre-potent reward seeking to wait for a larger reward (Capuzzo and Floresco, 2020; Cools, 2008). Research suggests that the expression of similar behavioral actions (including inhibition) is dependent on the motivational context (aversive vs. appetitive), which influences the likelihood of selecting a specific action in a specific motivational context (Wang and Delgado, 2021). However, in neuropsychiatric research, depressive disorders are often studied with regard to reward-seeking contexts, and anxiety disorders with regard to avoidance contexts, with little emphasis on the other motivational context. Symptoms of anxiety and depression have been associated with avoidance, typically operationalized via active avoidance and via questionnaires, as threats are overestimated (Bishop and Gagne, 2018; Browning, Behrens, Jocham, O’Reilly, and Bishop, 2015; Cléry-Melin, Schmidt, Lafargue, Baup, Fossati, and Pessiglione, 2011; Mkrtchian, Aylward, Dayan, Roiser, and Robinson, 2017; Ottenbreit, Dobson, and Quigley, 2014). In depression, reward-seeking may also be impaired due to a lack of motivation to obtain rewards (Alloy, Olino, Freed, and Nusslock, 2016; Bishop and Gagne, 2018). Past research has established the importance of avoidance and reward-seeking behaviours in helping us navigate our environment and stay safe (Krypotos, Effting, Kindt, and Beckers, 2015; LeDoux, Moscarello, Sears, and Campese, 2017). However, active vs. inhibitory subtypes of these behaviours have not typically been distinguished – especially through objective measures of observable behavior.

Gender - as a culturally defined construct - may also be an important variable in this relationship. For example, gender differences have been identified in the presentation and incidence of mood and anxiety disorders, such that women have higher rates of depression and present more often with depression than men (Altemus, Sarvaiya, and Neill Epperson, 2014; Kessler, 2006; Parker and Brotchie, 2010) and have rates of anxiety disorders that are twice as high as those of men (McLean, Asnaani, Litz, and Hofmann, 2011; Pittig, Treanor, LeBeau, and Craske, 2018). However, we do not know how these gender differences manifest themselves in avoidance or reward-seeking behaviours. Although mood and anxiety disorders are often comorbid, they also manifest with distinct symptoms and courses that would require distinct strategies to treat in a clinical context (Goldstein-Piekarski, Williams, and Humphreys, 2016; McLean, Asnaani, Litz, and Hofmann, 2011). In the present study, we ask how indices of anxiety and depression levels impact active vs. inhibitory avoidance and reward-seeking behaviours in a community population of young adults with a wide range of depressive and anxiety scores ranging from minimal to severe.

### The role of mood disorder symptoms and gender differences in avoidance and reward-seeking

It has been proposed that mood and anxiety disorder symptoms shift the perceived value and costs of avoidance and reward-seeking in sub-optimal ways. The Altered Computations Underlying Decision Making (ACDM) framework (Bishop and Gagne, 2018) proposes that anxiety is linked to underestimation of the effort cost in avoiding an aversive outcome and that depression is linked to overestimation of the effort cost in obtaining a reward. These effort costs interact with the perceived value of avoidance or reward-seeking to inform one’s decision on whether or not to engage in the behaviour. Past experimental work has also identified impairments in physical effort deployment for reward in populations with depression (Pessiglione, Vinckier, Bouret, Daunizeau, and Le Bouc, 2018; Treadway, Buckholtz, Schwartzman, Lambert, and Zald, 2009; see Culbreth, Moran, and Barch, 2018 for a review) and anxiety (Wang and Delgado, 2021). However, work linking mood and anxiety disorders to impairments in adaptive avoidance and reward-seeking often focuses on these avoidance and reward-seeking behaviours as unitary processes. As such, we still do not know how shifts in perceived effort costs linked to mood and anxiety disorders manifest themselves in active or inhibitory avoidance or reward-seeking.

To better understand the degree to which depressive and anxiety scores contribute to active and inhibitory forms of avoidance or reward-seeking a rigorous assessment of effort deployment in these behaviors is needed. People with Major Depressive Disorder (MDD) show a reduction in selecting high-effort, high-reward options on effort-based decision making tasks. This behaviour is potentially symptomatic of a larger-scale motivational deficit (Pessiglione, Vinckier, Bouret, Daunizeau, and Le Bouc, 2018; Treadway, Bossaller, Shelton, and Zald, 2012; Treadway, Buckholtz, Schwartzman, Lambert, and Zald, 2009). If maladaptive effort deployment is a primary characteristic of mood and anxiety disorders, then we might expect active avoidance and reward-seeking to be impaired more than inhibitory forms of these behaviours overall (Culbreth, Moran, and Barch, 2018). Anxiety, especially when co-occurring with high levels of depression, has also been shown to impair our sensitivity to rewards (Auerbach, Pagliaccio, Hubbard, Frosch, Kremens, Cosby, Jones, Siless, Lo, Henin, Hofmann, Gabrieli, Yendiki, Whitfield-Gabrieli, and Pizzagalli, 2022; Dillon, Rosso, Pechtel, Killgore, Rauch, and Pizzagalli, 2014); however, whether anxiety’s impact on reward-seeking differs for active or inhibitory behaviours is not yet clear.

Additionally, individual differences in the presentation and severity of mood and anxiety disorders - beyond the mere presence or absence of the disorder - may manifest with different patterns of active vs. inhibitory behaviours depending on the motivational context. Among these patterns, gender differences are especially prominent. Women generally present with higher levels of depression (Parker and Brotchie, 2010) and experience depression comorbid with anxiety more often than men (Kessler, 2006; McLean, Asnaani, Litz, and Hofmann, 2011; Ottenbreit, Dobson, and Quigley, 2014). Thus, the impact of mood and anxiety disorders on our ability to avoid aversive outcomes and seek out rewarding outcomes may be linked to gender differences that affect the motivational deficits these disorders present. If gender differences - looking across a full range of depressive and anxiety scores captured on self-report scales (Beck, Steer, and Brown, 1996) - predict differences in performance in a gender-dependent manner, then our study may help elucidate how gender differences in depressive and anxiety scores translate to changes in real-life behaviour.

In order to bring our understanding of mood disorder symptoms into a framework that acknowledges differences in active vs. inhibitory avoidance and reward-seeking behaviours, we must consider both anxiety and depression in a framework that directly investigates their impact on these behaviours, and how depressive and anxiety symptoms might interact to impair effective avoidance and reward-seeking. While the relationships between anxiety and avoidance (Bishop and Gagne, 2018; Levita, Hoskin, and Champi, 2012; Norbury, Robbins, and Seymour, 2018), and depression and reward-seeking (Alloy, Olino, Freed, and Nusslock, 2016; Rizvi, Pizzagalli, Sproule, and Kennedy, 2016; Treadway, Bossaller, Shelton, and Zald, 2012) are well established, those between anxiety and reward-seeking, as well as depression and avoidance, have yet to be fully characterized.

### An effortful avoidance and reward-seeking study

Despite established gender differences in the prevalence and presentation of mood disorder symptoms (Kessler, 2006; Parker and Brotchie, 2010; Thompson and Bland, 2018), it is not known how the relationship between mood and anxiety disorder symptom levels, and avoidance and reward-seeking, differs by gender. Gender differences in motivational deficits may lead to unique patterns in active and inhibitory behaviours, but this has not been examined either. As such, in this work, we ask: 1) whether anxiety and depression symptoms predict accuracy and effort deployment in active/inhibitory avoidance vs. reward-seeking; and 2) whether there are gender differences in the relationship between mood disorder scores and accuracy. We predicted that anxiety scores would significantly predict participants’ accuracy and the amount of effort they were able to deploy in avoidance behaviours (Bishop and Gagne, 2018) and that depression scores would significantly predict participants’ accuracy and effort in reward-seeking behaviours (Bishop and Gagne, 2018). We also predicted that the relationship between mood disorder scores – especially depression scores – and task performance (accuracy and effort) would be differ by gender, such that women would have lower performance than men in the task given higher depression scores (Parker and Brotchie, 2010).

To address these questions, the present study examined both avoidance and reward-seeking, each with two community online samples - undergraduates and online workers - with a broad distribution of mood disorder scores. Both studies were reverse-translated with modification from a series of rodent studies investigating deficits in active and inhibitory avoidance and reward-seeking behaviours (Capuzzo and Floresco, 2020; Piantadosi, Yeates, and Floresco, 2018). Our studies are the first to combine intermixed active and inhibitory avoidance (Levita, Hoskin, and Champi, 2012) or reward-seeking trials with increasing effort requirements throughout the task, requiring participants to switch between withholding physical effort on inhibitory trials and deploying increasing amounts of effort on active trials in each task. This design allows us to directly compare performance on active and inhibitory trials in the context of increasing effort demands. Increasing effort demands may also pull out differences in selecting between active vs. inhibitory strategies.

## Materials and Methods

### Participants

We powered each study to detect a moderate-sized main effect of *d* = 0.15 obtained with a previous study of *N* = 217 participants using the fabs R package (Biesanz, 2020), resulting in a target sample size of *N* = 549. Demographic information for all studies can be found in Table **??**. For each study, we collected data from two samples: an undergraduate population and an online worker population. The study was approved by the University of British Columbia Behavioural Research Ethics Board (BREB) under certificate H20-01388.

### Study 1 (Avoidance)

We recruited undergraduate participants at the University of British Columbia to participate online in our study. These participants received one percentage point towards their grade in a psychology course of their choosing for completing the study. Of these participants, *N* = 311 finished the study, of which *N* = 39 were excluded for not completing the pre-task survey, having below 50% accuracy on active or inhibitory avoidance trials, spending over 100 s on any given attention check, or not responding to all Beck Anxiety Inventory (BAI) questions. As such, data from *N* = 272 participants was used in the data analysis.

Additionally, we recruited paid online workers from around the world (*N* = 310) on the Prolific online study platform (https://www.prolif ic.co/). These participants received GBP £8.07 for completing the study. Of these participants, *N* = 294 finished the study, of which *N* = 22 were excluded for not completing the pre-task survey, having below 50% accuracy on active or inhibitory avoidance trials, spending over 100 s on any given attention check, or not responding to all Beck Anxiety Inventory (BAI) questions. As such, data from *N* = 272 participants was used in the data analysis.

### Study 2 (Reward-seeking)

We recruited undergraduate participants at the University of British Columbia to participate online in our study. These participants received one percentage point towards their grade in a psychology course of their choosing and a CAD $5.00 gift card from Starbucks for completing the study. Of these participants, *N* = 83 finished the study, of which *N* = 43 were part of a separate task condition with visual appetitive stimuli that is beyond the scope of this paper and *N* = 4 were excluded for not completing the pre-task survey, having below 50% accuracy on active or inhibitory avoidance trials, spending over 100 s on any given attention check, or incorrectly responding to a pre-task attention check. As such, data from *N* = 36 participants was used in the data analysis.

Additionally, we recruited paid online workers from around the world (*N* = 309) on the Prolific online study platform. These participants received GBP £8.07 and a £2.69 bonus for completing the study. Of these participants, *N* = 300 finished the study, of which *N* = 26 were excluded for not completing the pre-task survey, having below 50% accuracy on active or inhibitory avoidance trials, spending over 100 s on any given attention check, or incorrectly responding to a pre-task attention check. As such, data from *N* = 274 participants was used in the data analysis.

Overall, the excluded sample across both studies was 29.17% female and 70.83% male, while the analyzed sample was 45.90% female and 54.10% male.

## Materials and Methods

### Stimulus presentation

We used PsychoPy 2020.1.2 (RRID: SCR_006571) via the Pavlovia online study platform (Peirce, Gray, Simpson, MacAskill, Höchenberger, Sogo, Kastman, and Lindeløv, 2019). Participants completed the study online on their own computers; they were not allowed to complete the study on mobile devices or tablets.

### Stimuli

Cues indicating active or inhibitory trials were dark blue squares and circles with a thin black border and were generated by PsychoPy 2020.1.2 (Peirce, Gray, Simpson, MacAskill, Höchenberger, Sogo, Kastman, and Lindeløv, 2019) (Fig. **??**); they subtended a visual angle of about 11.5*^◦^* x 11.5*^◦^*. All stimuli were presented against a grey background (RGB value [0,0,0] on a scale from -1 to 1). If participants responded incorrectly on any trial in the avoidance studies, an aversive sound was played for 2000 ms. The aversive sounds were randomly selected from a set of eight screeching and scraping sounds created by our lab and ranked as highly aversive by four independent raters and in a pilot study.

Participants completed a series of questionnaires before beginning the main task. These were the State-Trait Anxiety Inventory, form Y-2 (STAI Y-2) (Spielberger, 2008); the Beck Depression Inventory II (BDI) (Beck, Steer, and Brown, 1996); the Beck Anxiety Inventory (BAI) (Steer and Beck, 1997); the Behavioral Activation for Depression Scale (BADS) (Kanter, Mulick, Busch, Berlin, and Martell, 2007); the Generalized Anxiety Disorder Scale (GAD-7) (Spitzer, Kroenke, Williams, and Löwe, 2006); and the Behavioural Inhibition Scale and Behavioural Activation Scale (BIS/BAS) (Carver and White, 1994). In our data analysis, we looked at results from the BDI and BAI as these clinically validated scales most directly capture participants’ levels of current depressive and anxiety scores. The BADS, GAD-7, and BIS/BAS capture specific behavioural facets of depression and anxiety that are less relevant to understanding overall effects of mood and anxiety disorders on avoidance and reward-seeking and were not analyzed in this study. We used the BAI as our primary measure of anxiety scores as it is the most widely used and validated among the anxiety scales we included (Fydrich, Dowdall, and Chambless, 1992) and as its structure parallels that of the BDI.

### Procedure

#### Avoidance task

A graphical overview of the avoidance task is provided in Fig. **??**A.

After an introduction screen, participants completed an effort calibration to control for differences in baseline effort ability and keyboard sensitivity. They were instructed to press the spacebar on their computer as many times as possible within a five-second period when a thermometer appeared on screen. Each time they pressed the spacebar, the thermometer would increase in height in order to incentivize participants to press the spacebar as many times as possible. Afterwards, participants repeated this effort calibration. This second calibration was identical to the first except that the thermometer would increase by only half the amount per press that it did for the first calibration, in order to incentivize participants to press more times during the second calibration and thereby better capture the participant’s maximum effort capability.

Following the effort calibration, participants completed an audio calibration to control for differences in audio cards and speakers. Here, participants were presented with a series of three one-second 2400 Hz sine tones - spaced by one second - at volumes of -50 dB, -30 dB, and -10 dB from maximum. After listening to these tones, participants were asked whether the first tone was barely heard and the final tone was aversive but not painful (Neumann and Waters, 2006). If this was not the case, participants were asked to adjust the volume on their computer and play the three tones again, repeating the process until the sound met these criteria - equalizing the experience of the sounds across participants. This computer volume was then used for the rest of the task.

After calibrating their physical effort capability and the volume of the aversive stimuli in the task, participants read instructions indicating the shape to which they would have to respond with multiple spacebar presses as well as the shape to which they would have to withhold their response. They also heard an example of the aversive sound that would be played if they made an incorrect response during the task.

In order to gain exposure to the stimuli and task contingencies, participants completed a series of practice trials (Fig. **??**A). This consisted of five trials in which participants had to make an active avoidance response - pressing the spacebar several times to avoid hearing an unpleasant sound; five trials in which participants had to make an inhibitory avoidance response - not pressing the spacebar to avoid hearing an aversive sound; and ten trials that intermixed these active and inhibitory trials.

On each trial, participants first viewed a grey screen with a white fixation cross for a mean duration of 2000 ms with a standard deviation (SD) of 1200 ms, jittered according to a normal distribution with these parameters on each trial. Participants then saw a visual cue - either a blue circle or a blue square - for 2000 ms. The cues used for active and inhibitory trials were pseudorandomly intermixed between participants. While this cue was on-screen, participants had to press the spacebar multiple times on active avoidance trials or withhold pressing on inhibitory avoidance trials. On active trials, the number of presses required was set according to the average number of presses made during the two effort calibration trials, such that participants who pressed fewer times during the calibration would have to press fewer times to achieve criterion during the task. The initial criterion was 5 presses given an average of 18 or fewer presses during calibration; a criterion of 6 presses given an average of 19-33 presses inclusive during calibration; and 7 presses given an average of 34 or more presses during calibration.

If participants made an incorrect response (pressing an insufficient number of times on active trials or pressing at all on inhibitory trials), participants heard an aversive sound and saw a fixation cross for 2000 ms. This aversive sound was taken from a set of ten sounds created by our lab and rated as highly aversive. All sounds were scraping sounds that had unpleasant psychoacoustic properties shown to reliably induce aversive responses (Neumann and Waters, 2006) at a variety of frequencies. If participants made a correct decision (pressing a sufficient number of times on active trials or not pressing on inhibitory trials), they saw a fixation cross surrounded by a white border that acted as a safety signal on the edges of the screen for 500 ms.

After completing the practice trials and viewing a final screen reminding them of the instructions, participants began the main task. This consisted of up to 168 active avoidance trials and 72 inhibitory avoidance trials (70% active and 30% inhibitory), pseudorandomized such that no more than 6 active trials or 3 inhibitory trials appeared in a row. On the 15th trial and every 40 trials thereafter, an attention check appeared asking participants to press a key corresponding to the letter they heard, to ensure that they were attending to the task and able to hear auditory stimuli. Every 20 trials, the number of button presses required on active trials increased by one press - this increased the effort demands on active trials across the task. The task continued until the participant responded correctly on half or less than half of the last 20 active trials - at this point, the breakpoint was reached and the participant was thanked for completing the task.

#### Reward-seeking task

A graphical overview of the reward-seeking task is provided in Fig. **??**B.

The design of the reward-seeking task was identical to that of the avoidance task, with the following exceptions. First, the practice blocks were based on criterion-based advancement in order to increase consistency with the design of other reward-seeking studies in our lab. Participants had to achieve at least 80% accuracy in each of the active, inhibitory, and intermixed reward-seeking trial blocks in order to advance; each block would repeat until they achieved each criterion. Second, if the participant made a correct decision during a trial, they would see a screen indicating that they had gained 5 points along with a sum of their points thus far; if the participant made an incorrect response during a trial, they would see a screen indicating that they had gained 0 points along with a sum of their points thus far. Both screens appeared for 1500 ms. Undergraduate participants received a CAD $5 gift card as a reward in addition to course credit for completing the task; online workers received a GBP £2.69 payment as a reward in addition to their payment for completing the task. Finally, as this task did not incorporate audio, no volume check or audio-based attention check was included.

### Data analysis

All data was analyzed using R 4.1.1 “Kick Things” (R Development Core Team, 2011) through RStudio (Booth et al., 2018). On each behavioural task, we measured: 1) participants’ sensitivity (*d′*), operationalized as the rate of correct active trials (hit rate) minus the rate of incorrect inhibitory trials (false alarm rate) and calculated using the dprime function in the psycho R package (Makowski, 2018) as the Z value of the hit rate minus the Z value of the false alarm rate; 2) effort on each trial type, operationalized as the number of presses made relative to criterion on each trial, averaged per block. 3) participants’ depressive and anxiety scores, operationalized as their BDI (Beck, Steer, and Brown, 1996) and BAI (Steer and Beck, 1997) scores respectively; and 4) breakpoint, operationalized as the trial number on which the participant responded incorrectly on half or less than half of the last 20 active trials. Breakpoint captures the point at which the effort demands of the task are no longer attainable for the participant, which is impaired in people with depression (Hershenberg, Satterthwaite, Daldal, Katchmar, Moore, Kable, and Wolf, 2016). On inhibitory trials, effort reflects the total number of presses made, to capture mistakes in which responses were still made on inhibitory trials. In order to capture all task parameters parsimoniously in our analyses, we analyzed sensitivity for avoidance and reward-seeking through four linear models in which either anxiety scores (BAI scores) or depression scores (BDI scores), together with gender and sample (university undergraduates vs. online workers), were included as predictors. We also analyzed effort for avoidance and reward-seeking through four multi-level models (using the lmerTest R package (Kuznetsova, Brockhoff, and Christensen, 2017)) in which either anxiety scores (BAI scores) or depression scores (BDI scores), together with gender, sample, and block of 28 active trials, were included as fixed effects and participant was a random effect. Lastly, we analyzed breakpoint for avoidance and reward-seeking using two linear models in which either anxiety scores (BAI scores) or depression scores (BDI scores), together with gender and sample (university undergraduates vs. online workers), were included as predictors. Linear models were used for sensitivity and breakpoint as these factors did not differ within participants, unlike effort - which differed between active and inhibitory trials, making a multi-level model appropriate. All analyses were Bonferroni corrected for multiple comparisons. All confidence intervals were based on 1000 bootstrap replications using the confintr R package (Mayer, 2022). All BDI and BAI scores were divided by the maximum number of points possible on each scale to obtain proportion scores. This was necessary because of the exclusion from our surveys of a BDI question relating to suicidality as the inclusion of such a question was not covered in our ethics, and because of the omission from our surveys of a BAI question that was used as an attention check – rendering raw BDI and BAI scores not comparable to those of other studies.

## Results

### Demographics

Participant’s reported gender and sex heavily overlapped. Of those reporting their gender as female, there was a 97.53% overlap with reported sex in women and 97.67% overlap in men on the avoidance tasks. There was a 98.84% overlap with reported sex women and 97.80% overlap in men on the reward-seeking task. For this reason, the following results are expressed in terms of gender only. Women reported higher levels of depressive scores (*t*(720.46) = -3.83, *p* < .001, *d* = 0.27) and anxiety scores (*t*(703.99) = -4.68, *p* < .001, *d* = 0.34) than men across samples (Table **??**, Figs. **??**, **??**). Across both studies, 20.98% of women and 15.11% of men were on medication for depression, and 19.35% of women and 14.89% of men were on medication for anxiety. For participants on these medications, BAI and BDI scores reflect their anxiety and depression scores in a medicated state and participants’ medication status was not included as a statistical control in our analyses. There were no significant differences in depression (*t*(582.52) = -1.61, *p* = 0.109, *d* = -0.12) or anxiety (*t*(588.90) = -0.22, *p* = 0.823, *d* = -0.02) between samples.

### Avoidance task

To account for participants’ bias to engage in active relative to inhibitory avoidance in general, we first calculated sensitivity (*d′*). Sensitivity reflects participants’ ability to correctly distinguish between active and inhibitory trials and deploy the required amount of effort on active trials only, while withholding effort on inhibitory trials. We additionally present results of active and inhibitory avoidance accuracy analyses at the link to the OSF repository in the data and code availability section (https://osf.io/2rd3f /). As all variance in sensitivity was between subjects, we ran a linear model analysis (Table **??**) to evaluate whether *d′* could be predicted from anxiety scores (BAI scores), gender, and sample in avoidance. Gender in interaction with anxiety scores significantly predicted sensitivity in the avoidance task such that women had lower performance with higher levels of anxiety (Fig. **??**A). There was also a BAI x Sample interaction such that participants with high anxiety in the global sample had lower performance than those with high anxiety in the undergraduate sample. We ran an additional linear model analysis (Table **??**) to evaluate whether *d′* could be predicted from depression scores (BDI scores), gender, and sample in avoidance (Fig. **??**B). Depression scores and sample significantly predicted *d′*, as did interactions between depression scores and gender, and between depression scores and sample - such that there was a significant interaction with women having lower sensitivity given higher depression scores. As with anxiety, women had lower performance with higher levels of depression. Women in the online worker sample had lower performance than those in the undergraduate sample given higher levels of depression.

Additionally, we explored the extent to which the amount of effort that participants exerted to avoid aversive outcomes changed across the avoidance task. As effort deployment could differ both between and within subjects, we conducted a multi-level model analysis (Table **??**) to evaluate whether effort deployment could be predicted from anxiety (BAI) scores, gender, sample, and block (28 active trials) in avoidance. This analysis revealed that participants deployed increasing amounts of effort during the task to meet increasing effort requirements (Fig. **??**A). There was also an interaction between gender and block qualified by a 3-way interaction between gender, BAI score, and block, indicating that the decrease in effort over time was associated with increased anxiety primarily in women (Fig. **??**B). We ran an additional multi-level model analysis (Table **??**) to evaluate whether effort could be predicted from depression (BDI) scores, gender, sample, and block in avoidance. Changes in effort across blocks during the task interacted with participants’ BDI scores and with gender, such that women with higher levels of depression deployed less effort relative to criterion in active avoidance.

Last, we examined whether breakpoint could be predicted from anxiety scores (BAI scores) or depression scores (BDI scores), gender, and sample in avoidance using two linear models (Tables **??**, **??**). None of these factors significantly predicted breakpoint in avoidance.

### Reward-seeking task

To evaluate participants’ bias to engage in active relative to inhibitory reward-seeking, we again calculated sensitivity (*d′*). We ran a linear model analysis (Table **??**) to evaluate whether *d′* could be predicted from anxiety scores (BAI scores), gender, and sample in reward-seeking. Only gender significantly predicted sensitivity in the reward-seeking task (Fig. **??**). We ran an additional linear model analysis (Table **??**) to evaluate whether *d′* could be predicted from depression scores (BDI scores), gender, and sample in avoidance. None of these factors predicted sensitivity in reward-seeking.

Additionally, we explored the extent to which the amount of effort that participants deployed to obtain reward changed across the reward-seeking task. We ran a multi-level model analysis (Table **??**) to evaluate whether effort deployment could be predicted from anxiety scores (BAI scores), gender, and sample in reward-seeking. This analysis revealed that effort decreased relative to criterion as the reward-seeking task progressed (Fig. **??**A). There were also effects of gender, with men deploying overall more effort than women, and anxiety, such that those with higher levels of anxiety were less able to deploy effort relative to criterion. These were qualified by an interaction between gender and anxiety such that women increased effort and men decreased it with higher levels of anxiety (Fig. **??**B). There were also a number of differences in effort deployment between samples, which interacted with a number of other predictors. We ran an additional multi-level model analysis (Table **??**) to evaluate whether effort could be predicted from depression scores (BDI scores), gender, sample, and block in reward-seeking. Only task block and gender predicted effort when depression scores were included as a predictor, such that women with higher depression scores deployed less effort relative to criterion in active reward-seeking.

Last, we examined whether breakpoint could be predicted from anxiety scores (BAI scores) or depression scores (BDI scores), gender, and sample in reward-seeking using two linear models (Tables **??**, **??**). None of these factors significantly predicted breakpoint in reward-seeking.

## Discussion

### Summary

In the present study, we investigated effects of gender and anxiety and depression levels in active and inhibitory avoidance and reward-seeking behaviours in a community population. Compared to men, women showed overall higher levels of self-reported depression and anxiety. Gender differences in task performance were in opposite directions depending on whether the task demanded avoidance or reward-seeking. Women showed lower sensitivity (*d′*), a measure of ability to correctly respond to active *and* inhibitory task demands, when avoiding an aversive outcome than men - but higher sensitivity when seeking reward. During active and inhibitory *avoidance*, gender interacted with anxiety level such that, in women, higher anxiety scores predicted lower sensitivity. There were also gender differences in effort deployment. Overall, throughout active *reward-seeking* trials, men made more effortful responses than women. Yet this finding was qualified by an interaction between gender and anxiety (over time) in both avoidance *and* reward-seeking, such that higher effort levels were associated with anxiety levels for women in both tasks, particularly early in the task. It should be noted that, in effortful reward seeking, effort deployment above criterion differed by sample such that online workers deployed more effort across the task and with increasing anxiety scores than undergraduate participants. Our findings illuminate gender differences in performance when both active and inhibitory responses are required for avoidance and reward-seeking in a context where effort is required to obtain the desired outcome. They also point to important boundary conditions for correlational effects that may vary between populations.

### Interpretation of results

#### Effects of gender and anxiety on avoidance task performance

We observed that, in avoidance, gender differences in performance (sensitivity; *d′*) where men had higher performance than women were moderated by anxiety, where women with higher anxiety had lower performance in the avoidance task. This finding is consistent with previous findings that avoidance behaviours are influenced by mood and anxiety levels (Pittig, Treanor, LeBeau, and Craske, 2018) which may partly explain overall gender differences in this task because of baseline gender differences in the prevalence of depression and anxiety (Parker and Brotchie, 2010). In particular, the observed gender differences in selecting the correct response on active vs. inhibitory trials (sensitivity; *d′*) when avoiding an unpleasant outcome could reflect more general gender differences in stress tolerance. In our study, women reported overall higher anxiety scores than men. Parker and Brotchie (2010) argued that women have a higher predisposition (diathesis) to stress than men and, in a context where an aversive outcome must be avoided, people with higher levels of anxiety scores may have a stronger impulse to act to avoid an aversive outcome (Bishop and Gagne, 2018). Thus, inhibiting a response may be especially difficult given a combination of high anxiety scores and a decreased tolerance for stress in women compared to men. Such a gender difference in diathesis could drive a reduced ability to inhibit effort when needed - an impairment in shifting from an active to an inhibitory strategy given higher levels of anxiety scores (Gustavson, Altamirano, Johnson, Whisman, and Miyake, 2017). Response inhibition can also require cognitive effort. People may have differing tendencies to make physically effortful responses to avoid aversive outcomes or obtain rewards, and to maintain an awareness that a cue is associated with withholding a response. Mood disorders have been associated with undervaluation of the reward of a cognitively effortful outcome (Grahek, Musslick, and Shenhav, 2020). Most of these findings have been associated with depression and reward-seeking. However, the reduced sensitivity with higher BAI scores observed in women in our avoidance task could reflect not only reduced motivation for physical effort on active trials with higher levels of anxiety but also reduced motivation to deploy cognitive effort to inhibit one’s response on inhibitory trials.

Gender differences in avoidance were also moderated by depression scores, such that women with higher depression scores performed worse on the avoidance task. Effects of depression scores, as measured by BDI scores, on task performance mostly reflected effects of anxiety. This is consistent with past findings that negative effects of anxiety and depression on motivated behaviours like avoidance and reward-seeking are often similar (Ottenbreit, Dobson, and Quigley, 2014), emphasizing the importance of a transdiagnostic approach when evaluating the impact of anxiety and depression on these behaviours (Culbreth, Moran, and Barch, 2018). It may also reflect the extent to which the BAI and BDI measure overlapping constructs, as reflected by the high correlation between BAI and BDI scores we observed (*t*(852) = 23.86, *p* = < .001, *r* = 0.63).

#### Effects of gender and anxiety on effort deployment in reward-seeking and avoidance

In reward-seeking, we observed an opposing relationship between gender and effort deployment to that in avoidance. Women deployed more effort relative to criterion at higher anxiety scores than men, while men continued to deploy less effort with higher anxiety scores. The diathesis effects that may impair effective effort deployment in avoidance may not be present in reward-seeking, as effects of an error are likely to be less stressful to participants. Therefore, the observed effort impairments in women may be valence-specific.

On active reward-seeking trials, men deployed more effort relative to criterion across the task than women. This could be caused by women having smaller wrists with which to generate physical force than men (Morse, Jung, Bashford, and Hallbeck, 2006), as well as men having - on average - higher levels of testosterone levels compared to women - which is associated with increased physical effort (Losecaat Vermeer, Riečanský, and Eisenegger, 2016) and risk tolerance (Cooper, Goings, Kim, and Wood, 2014). This initial difference in effort deployment capability is reflected in the finding that men pressed significantly more than women in the pre-task effort calibration across all tasks (*t*(814.99) = 7.02, *p* = < .001). Although our tasks did not have competitive elements, participants may still have completed the task with an eye towards maximizing performance. Since deploying more effort in the task would increase one’s chance of staying above criterion, this increased effort deployment in men could explain the increased active trial accuracy for men across the avoidance and reward-seeking tasks. It is important to qualify that gender has a significant cultural component, and cultural factors could also play a role in gender differences in effort deployment - perhaps via effects of a lower tolerance for stress on effort deployment (Parker and Brotchie, 2010). Importantly, in both tasks gender interacted with anxiety levels and, in the avoidance task, gender differences in effort was qualified by a relationship with anxiety early in the task, with women higher in anxiety scores deploying higher effort levels early on. Thus any gender differences in effort are complex, and vary with anxiety levels, motivational context, and likely other boundary conditions as well.

It should be noted that the relationship between higher anxiety scores and reduced avoidance sensitivity, while qualified by gender and sample, differs from previous predictions of improved avoidance in anxiety, such as those of Bishop and Gagne (2018). Bishop and Gagne framed this relationship in terms of active and not inhibitory avoidance, as they predicted that underestimations of effort cost would drive excessive avoidance behaviours. Anxiety scores may be associated with impairments to inhibitory avoidance precisely because of this bias towards action given the possibility of aversive outcomes, an effect that could be driven by a perceived lack of control over outcomes in the task (Wang and Delgado, 2021). Additionally, we did not observe a relationship between depression scores and accuracy or effort deployment in reward-seeking, as has previously been observed (Bishop and Gagne, 2018). As depression scores did not influence effort deployment, we can speculate that, in this task, the effort demands of the task did not deter those with higher depression scores from working for a reward.

Overall, participants deployed less effort in avoidance compared to reward-seeking; this could be a function of differences in motivation to engage in avoidance or reward-seeking. Motivation to complete the tasks can be driven in part by participants’ valuations of task-relevant stimuli (Bishop and Gagne, 2018). A major difference between our tasks arises in the outcome of an incorrect response. In avoidance, an incorrect response is associated with an aversive sound; in reward-seeking, it is associated with not receiving points. Although the salience of an aversive sound may suggest that it is more motivating and would therefore be associated with increased accuracy, hearing it may also be more demotivating - especially for participants with mood disorder symptoms. Hearing the aversive sound repeatedly could be a salient indicator of a lack of control over task outcomes (Wang and Delgado, 2021).

### Limitations

There are some limitations to our interpretation of our findings. First, since the dichotomy of the task demands is between effortful active trials and inhibitory trials that require no effort, we cannot compare the effects of high vs. low effort demands on inhibitory avoidance or reward-seeking behaviours. As such, our interpretation of the relationship between effort deployment and mood disorder symptoms only extends to active trials. Accuracy in the task was likely tied to participants’ effort capabilities, as increased effort deployment was required throughout the task on active trials to meet the criterion level of effort and make the correct response on the trial. However, we calibrated the criterion to participants’ effort ability and considered performance on inhibitory as well as active trials to reduce the reliance of task outcomes on individual differences in effort deployment. Additionally, as the proportion of active trials was greater than that of inhibitory trials, participants may have become increasingly fatigued on the majority of trials in the task. This fatigue from effort deployment, combined with boredom (from the task being repetitive) could be difficult to disentangle from other shifts in motivation to deploy effort throughout the task (e.g. those related to the value of avoidance or reward-seeking). However, as fatigue is likely to arise in most physically effortful tasks, our tasks still reflect real-world physical effort demands. Furthermore, as this study took place online, the study had to use repeated keyboard presses instead of other, more continuous or better-controlled measures of physical effort such as a grip squeeze (Aridan, Malecek, Poldrack, and Schonberg, 2019). However, repeated button presses have been validated as being physically effortful and have been used in in-person contexts (Gold, Strauss, Waltz, Robinson, Brown, and Frank, 2013).

When predicting avoidance sensitivity, we observed interactions between BAI and BDI scores and sample; when predicting reward-seeking effort, we observed an interaction between gender and sample. This suggests that performance differences related to anxiety, depression, or gender differ according to the demographic makeup of each sample. For example, women reported overall higher levels of anxiety than men. The interaction we observed between BAI scores and sample in predicting *d′* in the avoidance task may suggest a differing relationship between performance and anxiety levels between the younger, female-skewed online undergraduates and older, male-skewed online workers. As online workers were paid in money for participation while undergraduates received course credit, motivations for completing the task may also have differed between samples. In addition, the online workers were likely a more heterogeneous sample in terms of the devices and contexts in which they completed the task. It should also be noted that, whereas the avoidance study was well-balanced between males and females, the reward-seeking study had a substantially higher proportion of male to female participants - as a result of the higher proportion of males in the larger, online worker sample in this study. It may be that the interactions we observed between sample and gender in predicting reward-seeking effort deployment can be explained by this substantially higher proportion of male to female participants. However, the presence of significant effects that do not interact with sample in both studies suggest that sample differences did not explain a large proportion of the variance in our findings.

Additionally, with a final *N* of 310 participants’ data analyzed, the reward-seeking study fell short of our target sample size because of limitations in availability of the undergraduate sample. For this reason, it may have been underpowered to reliably detect higher-order interactions.

### Future work

Future studies could build on our findings by investigating how patterns of information about specific aspects of effortful avoidance and reward-seeking are instantiated in key brain regions. The posterior anterior cingulate cortex (pACC) and ventral striatum encode information about prospective gains given physical effort requirements (Aridan, Malecek, Poldrack, and Schonberg, 2019). These regions - and their homologues in rodents - have been shown to be differentially necessary for active vs. inhibitory avoidance (Piantadosi, Yeates, and Floresco, 2018) and reward-seeking (Capuzzo and Floresco, 2020). Investigating how these regions represent information on prospective threats and gains relative to effort costs could illuminate how we weigh the benefits and costs of deploying effort to obtain rewards and avoid aversive outcomes. Additionally, separating out different factors contributing to effort deployment through a computational modelling approach would be important to understand the individual contributions of various factors to participants’ performance. These factors could include action biases (Mkrtchian, Aylward, Dayan, Roiser, and Robinson, 2017), perceived value of avoidance or reward (Bishop and Gagne, 2018), or fatigue (Pessiglione, Vinckier, Bouret, Daunizeau, and Le Bouc, 2018). Furthermore, it would be helpful to evaluate whether subscales of mood disorder symptoms - potentially linked to subtypes such as anxious depression (Wurst, Schiele, Stonawski, Weiß, Nitschke, Hommers, Domschke, Herrmann, Pauli, Deckert, and Menke, 2021) - pull out factors that drive participants’ behaviours in avoidance and reward seeking. This analysis could further illuminate our observed gender differences - for example, to evaluate whether reduced avoidance sensitivity in women given increased anxiety scores is reflective of an anxious subtype of depression (Wurst, Schiele, Stonawski, Weiß, Nitschke, Hommers, Domschke, Herrmann, Pauli, Deckert, and Menke, 2021).

## Conclusion

Our studies address outstanding questions of whether a range of anxiety and depression scores predict performance (sensitivity) and effort deployment in avoidance and reward-seeking, and whether the relationship between performance and anxiety/depression levels is impacted by gender. We elicit both active and inhibitory avoidance and reward-seeking behaviours in a context that allows for direct comparisons between them, instead of considering avoidance and reward-seeking behaviours as unitary wholes. We highlight gender differences in each of these subtypes of avoidance and reward-seeking given varying levels of anxiety and depression scores, contextualizing past work on gender differences (Parker and Brotchie, 2010). In particular, we are the first to examine these proposed gender differences in an active and inhibitory avoidance and reward-seeking context. These findings could inform clinical interventions to address maladaptive deployment of avoidance and lack of motivation for reward-seeking, targeted by gender. Additionally, we link active avoidance and reward-seeking to motivation for physical effort deployment given varying levels of mood and anxiety disorder severity. As many tasks in life require physical effort deployment, understanding where it can be impaired is an important pursuit. Our findings underscore the importance of considering individual differences in the ways in which avoidance and reward-seeking can be impaired in life.

## Data and code availability

The data and materials for all experiments, as well as the code used to generate this manuscript and conduct all analyses, are available at https://osf.io/2rd3f /.

## Legend

**Figure 1:**
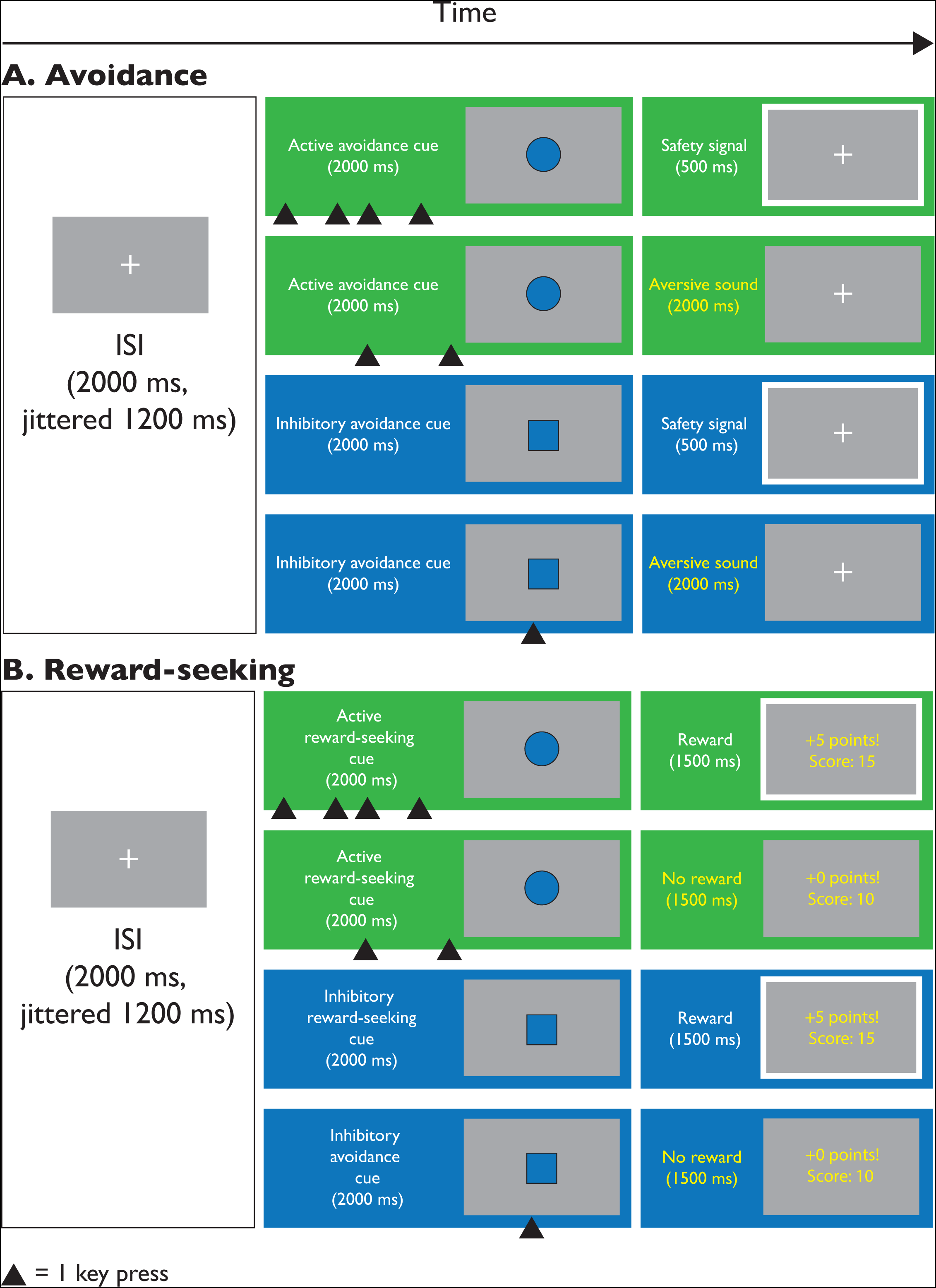
Trial layout diagram. A diagram of the active and inhibitory avoidance and reward-seeking tasks. In the avoidance task (A), after an inter-stimulus interval (ISI) with a fixation cross onscreen, participants were presented with a cue associated with active or inhibitory avoidance. For the active avoidance cue, participants had to respond with repeated spacebar presses to avoid hearing an aversive sound. For the inhibitory avoidance cue, participants had to withhold responding to avoid hearing an aversive sound. In the reward-seeking task (B), after the ISI, participants were presented with a cue associated with active or inhibitory reward-seeking. For the active reward-seeking cue, participants had to respond with repeated spacebar presses to obtain points towards a monetary reward. For the inhibitory reward-seeking cue, participants had to withhold responding to obtain points towards a monetary reward. ISI = Inter-stimulus interval.

**Figure 2:**
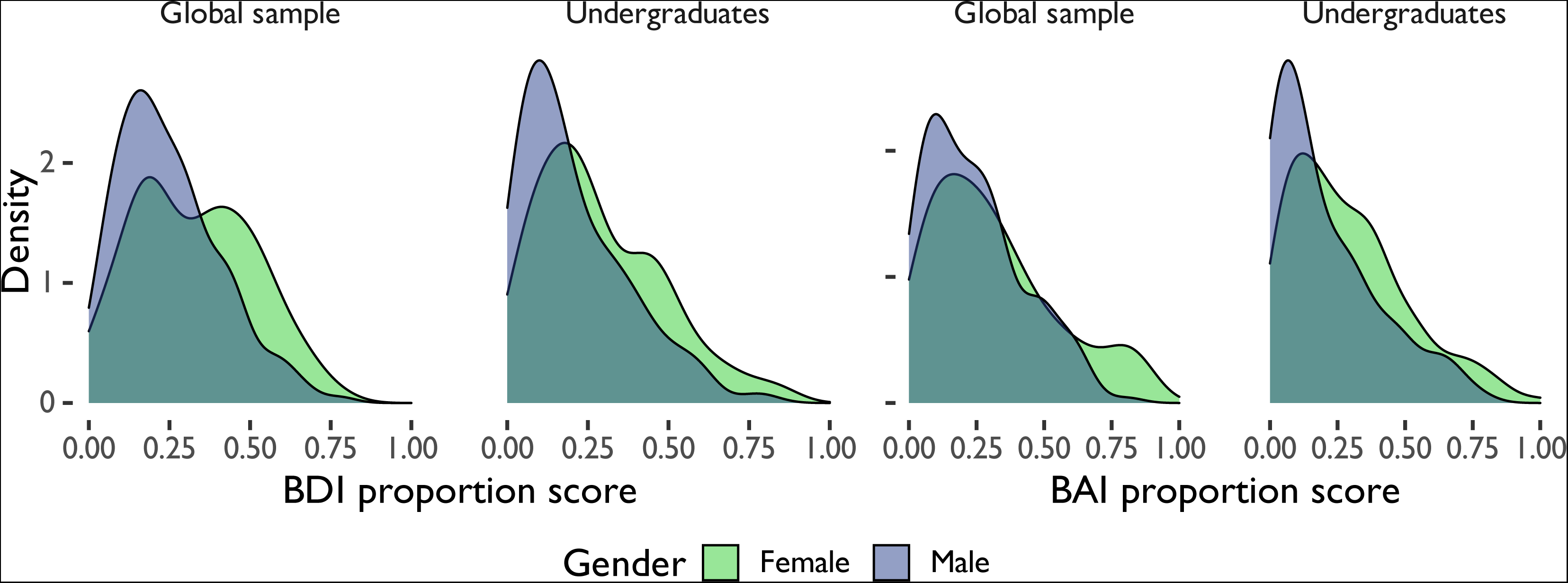
Avoidance demographics. Distribution of anxiety (BAI) and depression (BDI) scores by gender and sample. Proportion scores are scores divided by total possible score. BAI = Beck Anxiety Inventory, BDI = Beck Depression Inventory II.

**Figure 3:**
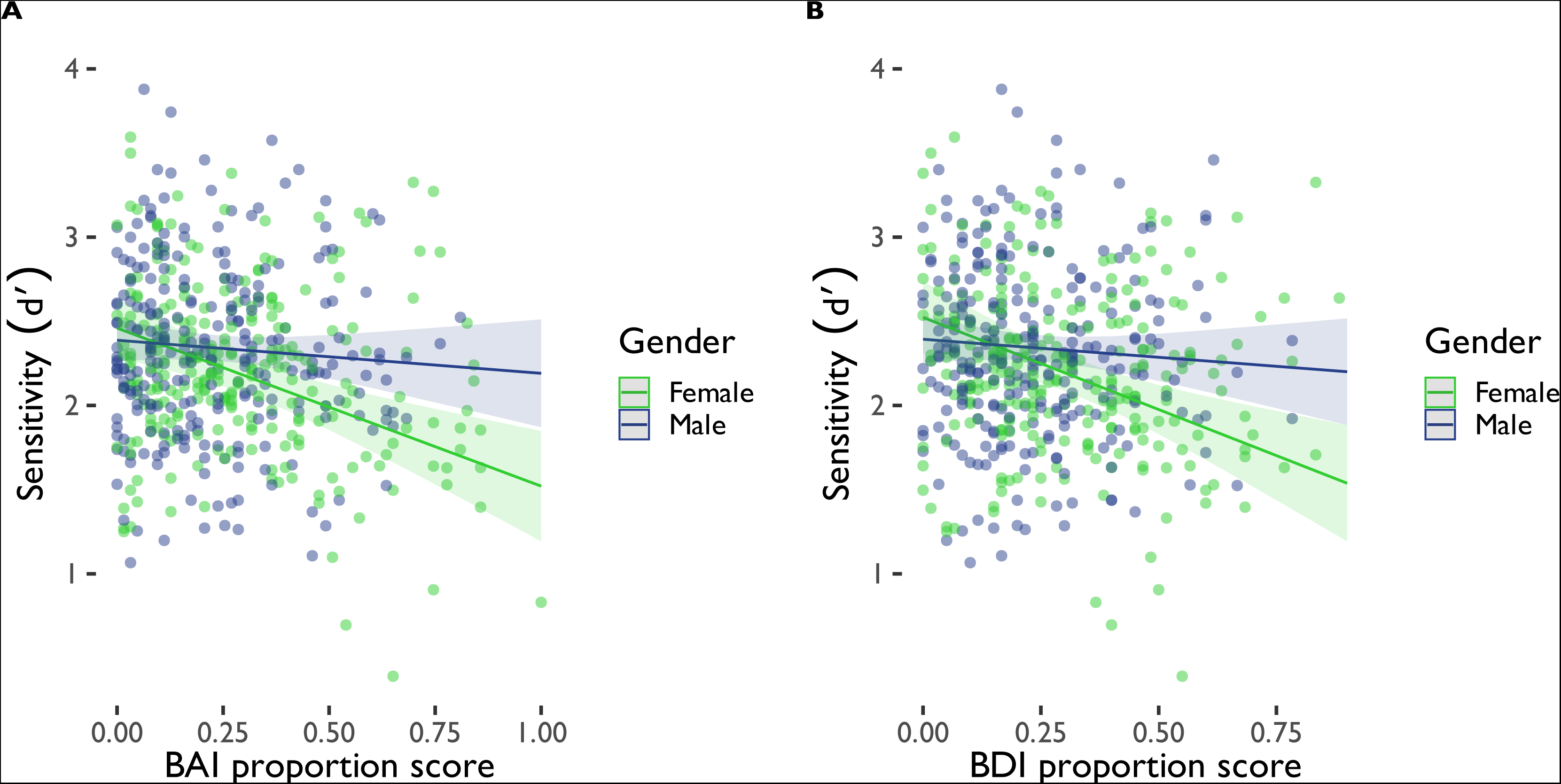
Linear model significant effects for sensitivity in avoidance. (A) Gender interacted with anxiety scores (BAI proportion scores) to explain sensitivity (*d′*) in the avoidance task. (B) Gender interacted with depression scores (BDI proportion scores) to explain sensitivity (*d′*) in the avoidance task. BAI = Beck Anxiety Inventory. BDI = Beck Depression Inventory II. Proportion scores are scores divided by total possible score.

**Figure 4:**
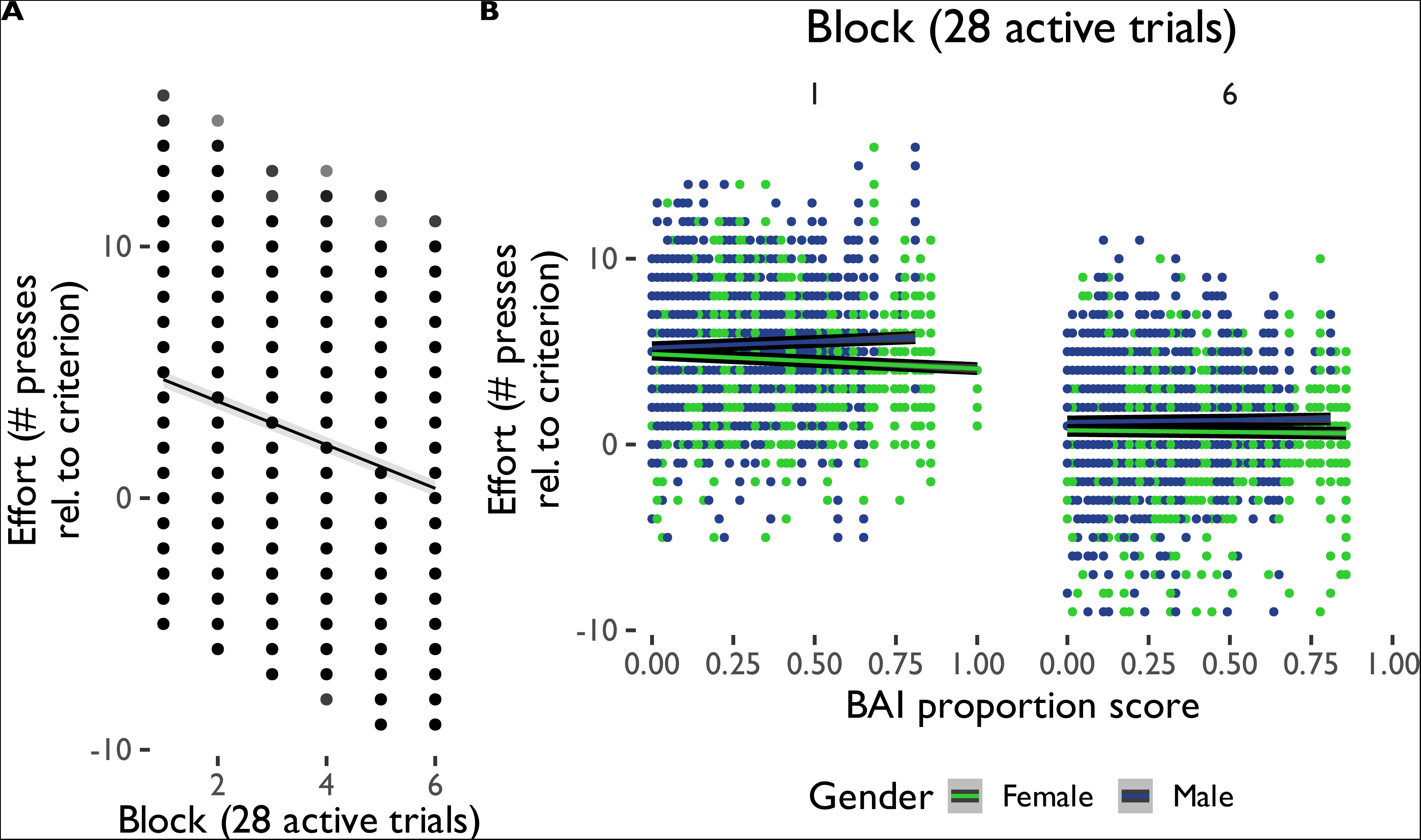
Avoidance effort deployment. (A) Multi-level model significant effects and interactions for effort in avoidance. Effort decreased relative to criterion as the avoidance task progressed, an effect that (B) interacted with anxiety scores (BAI) and gender. Proportion scores are scores divided by total possible score.

**Figure 5:**
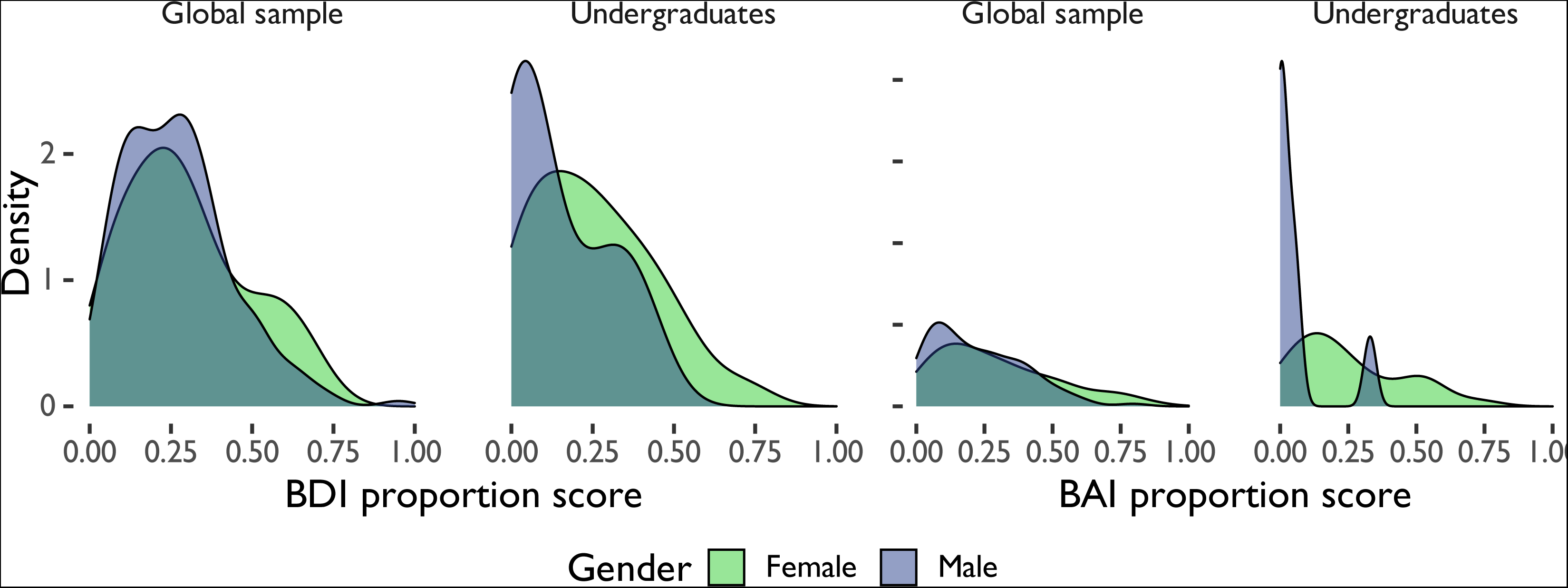
Reward-seeking demographics. Distribution of anxiety (BAI) and depressive (BDI) scores by gender and sample. Proportion scores are scores divided by total possible score. BAI = Beck Anxiety Inventory, BDI = Beck Depression Inventory II.

**Figure 6:**
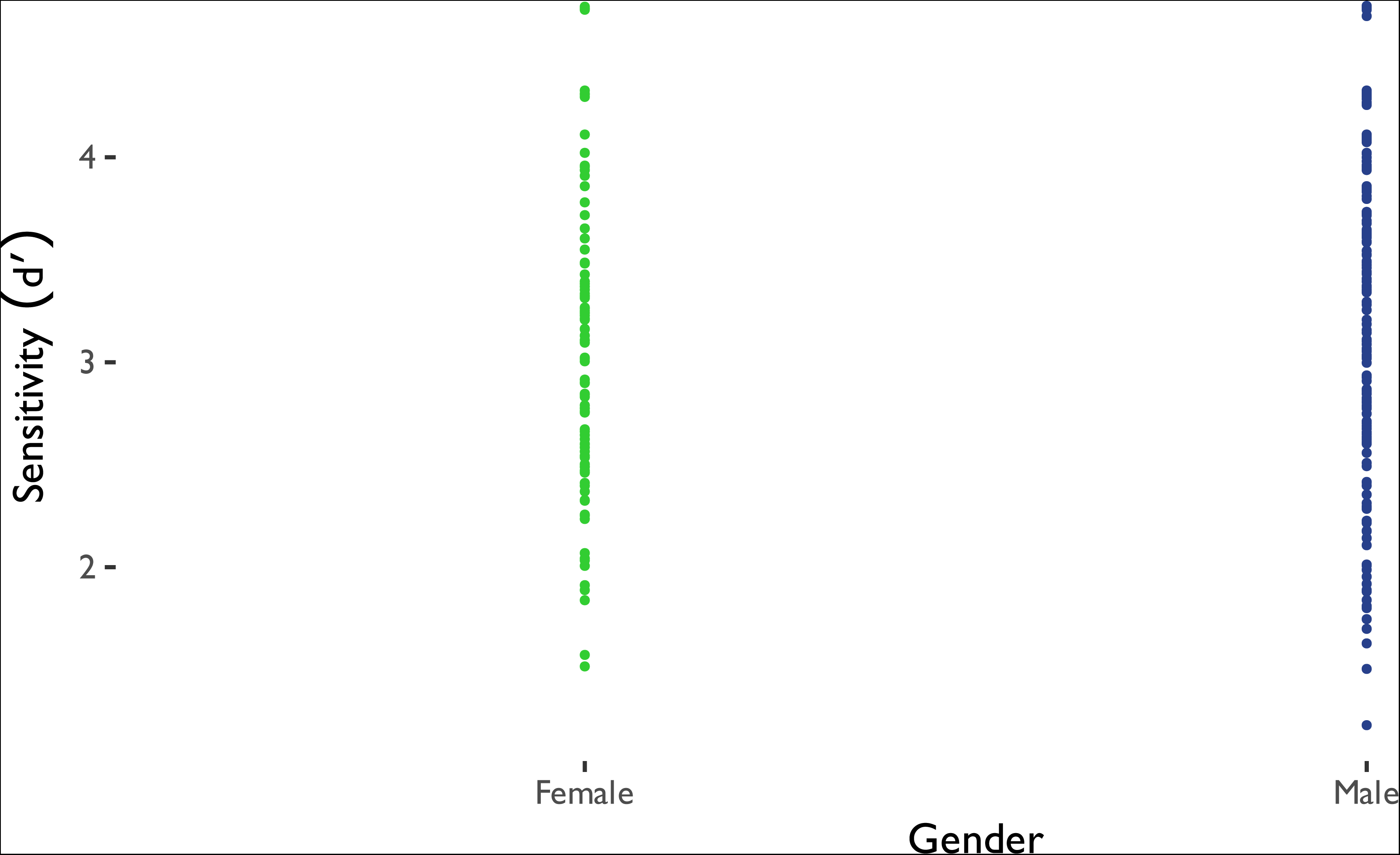
Linear model significant effects for sensitivity in reward-seeking. Gender explained sensitivity (*d′*) in the reward-seeking task. BAI = Beck Anxiety Inventory. Proportion scores are scores divided by total possible score.

**Figure 7:**
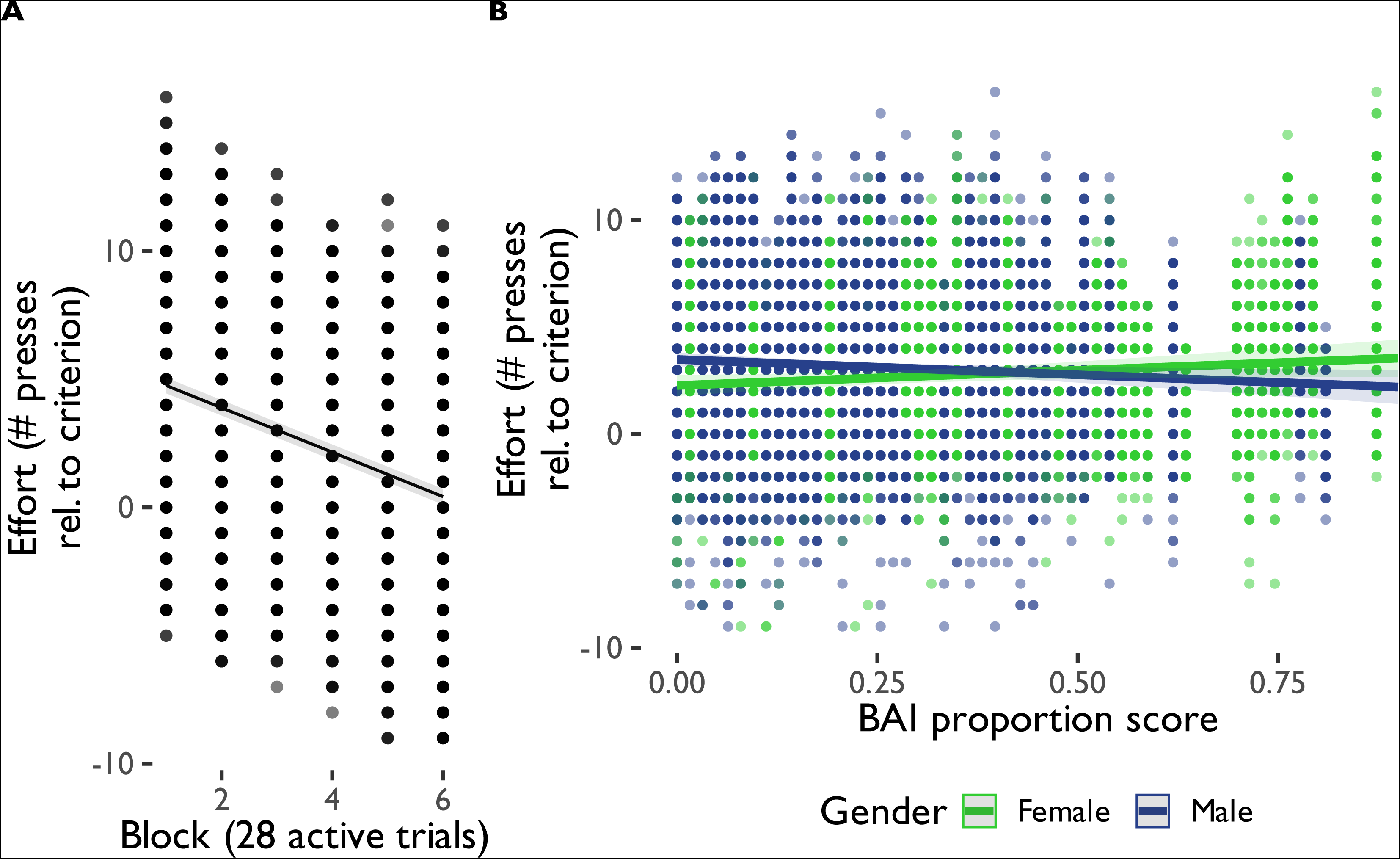
Reward-seeking effort deployment. (A) Multi-level model significant effects and interactions for effort in reward-seeking. Effort decreased relative to criterion as the reward-seeking task progressed, an effect that (B) interacted with anxiety scores (BAI) and gender. Proportion scores are scores divided by total possible score.

**Table 1:**
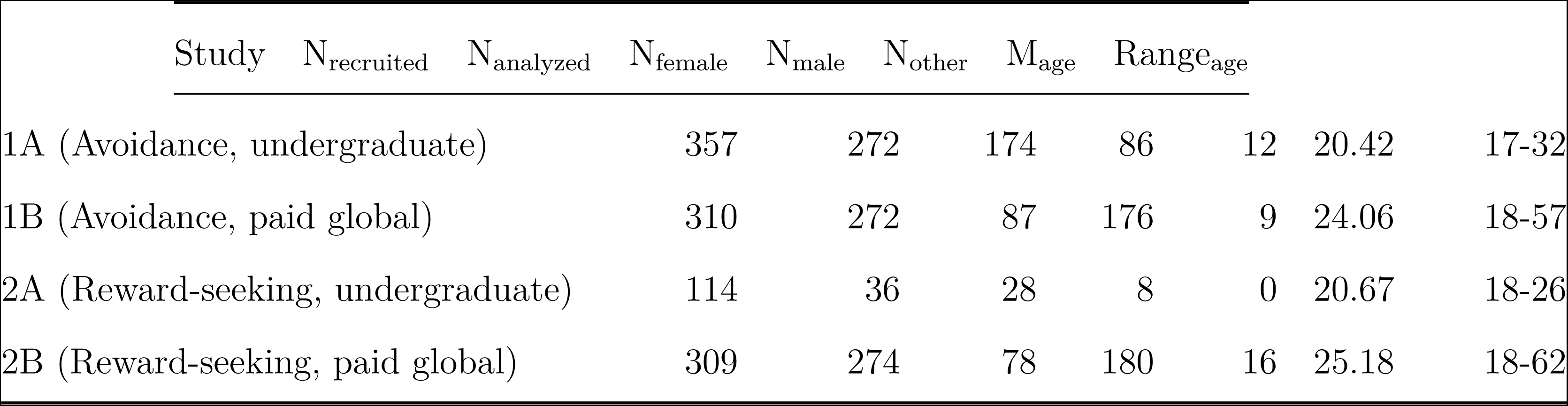
Demographic information for all participants.

**Table 2:** Mood disorder symptom statistics. Mean and SD Beck Depression Inventory II (BDI) and Beck Anxiety Inventory (BAI) proportion scores (score divided by total possible score).

| Task | Gender | M<sub>BDIprop</sub> | SD<sub>BDIprop</sub> | M<sub>BAIprop</sub> | SD<sub>BAIprop</sub> |
| --- | --- | --- | --- | --- | --- |
| Avoidance | Female | 0.30 | 0.19 | 0.29 | 0.22 |
|  | Male | 0.23 | 0.16 | 0.23 | 0.19 |
| Reward-seeking | Female | 0.29 | 0.19 | 0.28 | 0.21 |
|  | Male | 0.26 | 0.17 | 0.22 | 0.17 |

**Table 3:**
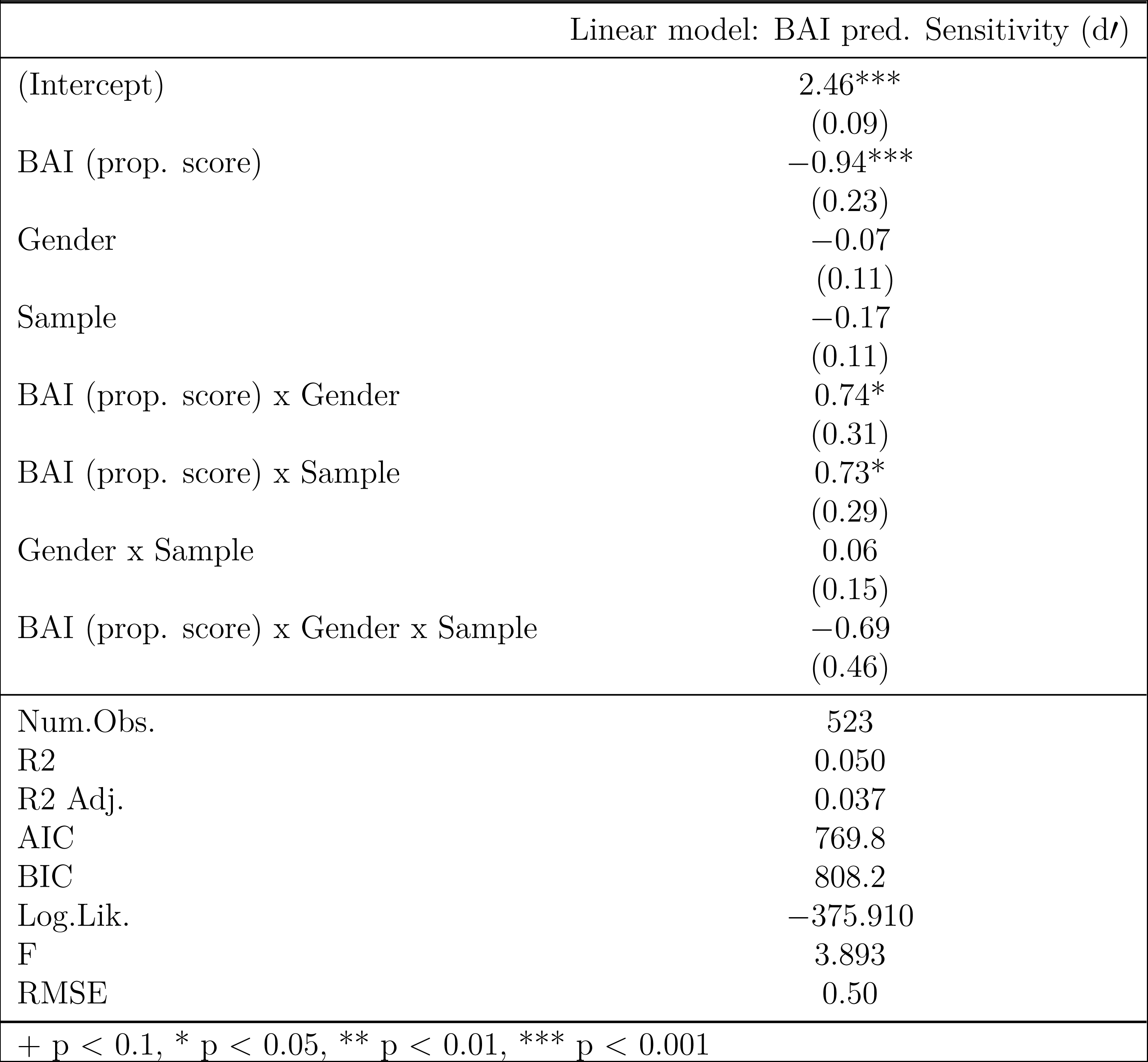
Linear model analysis coefficients and standard errors for sensitivity (*d′*) in avoidance. BAI (prop. score) = anxiety score on the Beck Anxiety Inventory. Proportion scores are scores divided by total possible score. AIC = Akaike information criterion, BIC = Bayesian information criterion, Log. Lik. = log likelihood, RMSE = root mean squared error.

**Table 4:** Linear model analysis coefficients and standard errors for sensitivity (d^t^) in avoidance. BDI (prop. score) = depression score on the Beck Depression Inventory II. Proportion scores are scores divided by total possible score. AIC = Akaike information criterion, BIC = Bayesian information criterion, Log. Lik. = log likelihood, RMSE = root mean squared error.

| | Linear model: BDI pred. Sensitivity ($d'$) |
| --- | --- |
| (Intercept) | 2.52***<br>(0.11) |
| BDI (prop. score) | -1.09***<br>(0.29) |
| Gender | -0.13<br>(0.13) |
| Sample | -0.29*<br>(0.13) |
| BDI (prop. score) x Gender | 0.88*<br>(0.38) |
| BDI (prop. score) x Sample | 1.11**<br>(0.35) |
| Gender x Sample | 0.16<br>(0.17) |
| BDI (prop. score) x Gender x Sample | -0.93+<br>(0.53) |
| Num.Obs. | 523 |
| R<sup>2</sup> | 0.042 |
| R<sup>2</sup> Adj. | 0.029 |
| AIC | 774.1 |
| BIC | 812.4 |
| Log.Lik. | -378.038 |
| F | 3.265 |
| RMSE | 0.50 |

+ $p < 0.1$ , \* $p < 0.05$ , \*\* $p < 0.01$ , \*\*\* $p < 0.001$

**Table 5:** Multi-level model analysis coefficients and standard errors for effort in avoidance. BAI (prop. score) = anxiety score on the Beck Anxiety Inventory. Proportion scores are scores divided by total possible score. AIC = Akaike information criterion, BIC = Bayesian information criterion, ICC = intraclass correlation, RMSE = root mean squared error.

|  | Multi-level model: BAI pred. Effort |
| --- | --- |
| (Intercept) | 5.74***<br>(0.26) |
| BAI (prop. score) | −0.63<br>(0.68) |
| Block | −0.91***<br>(0.02) |
| Gender | 0.29<br>(0.32) |
| Sample | −0.42<br>(0.33) |
| BAI (prop. score) x Block | 0.16***<br>(0.05) |
| BAI (prop. score) x Gender | 0.97<br>(0.92) |
| Block x Gender | 0.04*<br>(0.02) |
| BAI (prop. score) x Sample | −0.65<br>(0.87) |
| Block x Sample | −0.01<br>(0.03) |
| Gender x Sample | −0.19<br>(0.44) |
| BAI (prop. score) x Block x Gender | −0.19**<br>(0.06) |
| BAI (prop. score) x Block x Sample | −0.06<br>(0.07) |
| BAI (prop. score) x Gender x Sample | 0.94<br>(1.36) |
| Block x Gender x Sample | 0.04<br>(0.04) |
| BAI (prop. score) x Block x Gender x Sample | −0.08<br>(0.11) |
| SD (Intercept participant) | 1.42 |
| SD (Observations) | 1.91 |
| Num.Obs. | 52261 |
| R2 Marg. | 0.289 |
| R2 Cond. | 0.542 |
| AIC | 218248.5 |
| BIC | 218408.1 |
| ICC | 0.4 |
| RMSE | 1.90 |

**Table 6:** Multi-level model analysis coefficients and standard errors for effort in avoidance. BDI (prop. score) = depression score on the Beck Depression Inventory II. Proportion scores are scores divided by total possible score. AIC = Akaike information criterion, BIC = Bayesian information criterion, ICC = intraclass correlation, RMSE = root mean squared error.

|  | Multi-level model: BDI pred. Effort |
| --- | --- |
| (Intercept) | 5.67***<br>(0.32) |
| BDI (prop. score) | −0.41<br>(0.86) |
| Block | −0.92***<br>(0.02) |
| Gender | 0.24<br>(0.38) |
| Sample | −0.40<br>(0.38) |
| BDI (prop. score) x Block | 0.19**<br>(0.06) |
| BDI (prop. score) x Gender | 1.21<br>(1.11) |
| Block x Gender | 0.08**<br>(0.03) |
| BDI (prop. score) x Sample | −0.70<br>(1.04) |
| Block x Sample | 0.01<br>(0.03) |
| Gender x Sample | −0.31<br>(0.50) |
| BDI (prop. score) x Block x Gender | −0.34***<br>(0.08) |
| BDI (prop. score) x Block x Sample | −0.12<br>(0.08) |
| BDI (prop. score) x Gender x Sample | 1.54<br>(1.59) |
| Block x Gender x Sample | −0.01<br>(0.04) |
| BDI (prop. score) x Block x Gender x Sample | 0.12<br>(0.13) |
| SD (Intercept participant) | 1.42 |
| SD (Observations) | 1.91 |
| Num.Obs. | 52261 |
| R<sup>2</sup> Marg. | 0.289 |
| R<sup>2</sup> Cond. | 0.542 |
| AIC | 218241.9 |
| BIC | 218401.4 |
| ICC | 0.4 |
| RMSE | 1.90 |

**Table 7:** Linear model analysis coefficients and standard errors for breakpoint in avoidance. BAI (prop. score) = anxiety score on the Beck Anxiety Inventory. Proportion scores are scores divided by total possible score. AIC = Akaike information criterion, BIC = Bayesian information criterion, Log. Lik. = log likelihood, RMSE = root mean squared error.

|  | Linear model: BAI pred. Sensitivity (d') |
| --- | --- |
| (Intercept) | 133.41***<br>(7.78) |
| BAI (prop. score) | −18.55<br>(20.00) |
| Gender | 12.52<br>(9.52) |
| Sample | −5.39<br>(9.53) |
| BAI (prop. score) x Gender | 42.11<br>(27.05) |
| BAI (prop. score) x Sample | −12.37<br>(25.25) |
| Gender x Sample | 5.42<br>(12.91) |
| BAI (prop. score) x Gender x Sample | −30.85<br>(39.58) |
| Num.Obs. | 523 |
| R<sup>2</sup> | 0.105 |
| R<sup>2</sup> Adj. | 0.093 |
| AIC | 5440.4 |
| BIC | 5478.7 |
| Log.Lik. | −2711.198 |
| F | 8.668 |
| RMSE | 43.16 |

+ p < 0.1, \* p < 0.05, \*\* p < 0.01, \*\*\* p < 0.001

**Table 8:**
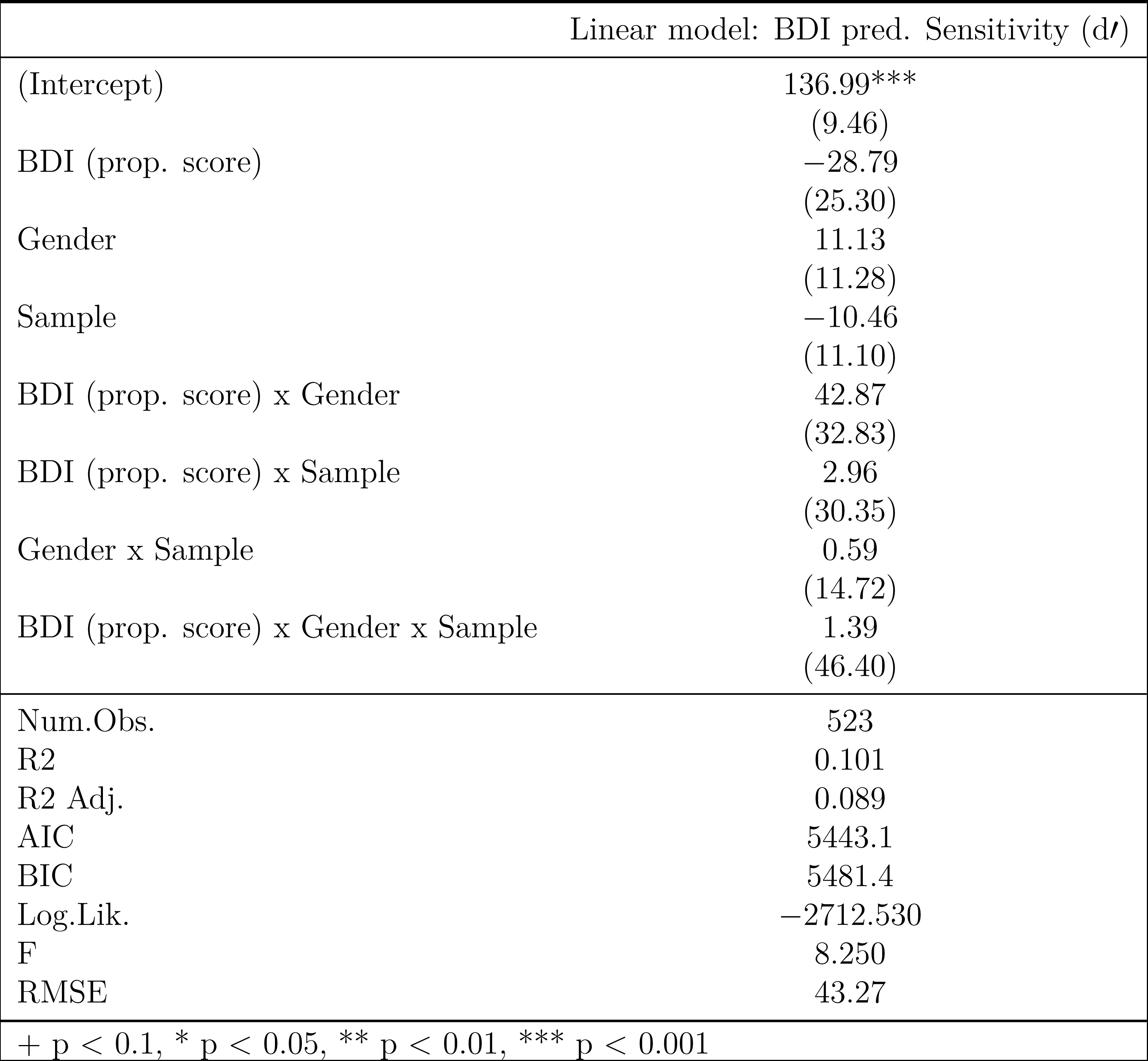
Linear model analysis coefficients and standard errors for breakpoint in avoidance. BDI (prop. score) = depression score on the Beck Depression Inventory II. Proportion scores are scores divided by total possible score. AIC = Akaike information criterion, BIC = Bayesian information criterion, Log. Lik. = log likelihood, RMSE = root mean squared error.

**Table 9:** Linear model analysis coefficients and standard errors for sensitivity (d^t^) in reward-seeking. BAI (prop. score) = anxiety score on the Beck Anxiety Inventory. Proportion scores are scores divided by total possible score. AIC = Akaike information criterion, BIC = Bayesian information criterion, Log. Lik. = log likelihood, RMSE = root mean squared error.

| | Linear model: BAI pred. Sensitivity ($d'$) |
| --- | --- |
| (Intercept) | 3.12***<br>(0.15) |
| BAI (prop. score) | −0.16<br>(0.40) |
| Gender | 0.34*<br>(0.17) |
| Sample | 0.15<br>(0.27) |
| BAI (prop. score) x Gender | −0.75<br>(0.52) |
| BAI (prop. score) x Sample | −0.48<br>(0.84) |
| Gender x Sample | −0.53<br>(0.42) |
| BAI (prop. score) x Gender x Sample | 3.16<br>(2.69) |
| Num.Obs. | 294 |
| R<sup>2</sup> | 0.041 |
| R<sup>2</sup> Adj. | 0.017 |
| AIC | 686.9 |
| BIC | 720.1 |
| Log.Lik. | −334.458 |
| F | 1.739 |
| RMSE | 0.75 |

+ $p < 0.1$ , \* $p < 0.05$ , \*\* $p < 0.01$ , \*\*\* $p < 0.001$

**Table 10:** Linear model analysis coefficients and standard errors for sensitivity (d^t^) in reward-seeking. BDI (prop. score) = anxiety score on the Beck Depression Inventory II. Proportion scores are scores divided by total possible score. AIC = Akaike information criterion, BIC = Bayesian information criterion, Log. Lik. = log likelihood, RMSE = root mean squared error.

| | Linear model: BDI pred. Sensitivity ($d'$) |
| --- | --- |
| (Intercept) | 3.09***<br>(0.16) |
| BDI (prop. score) | −0.05<br>(0.46) |
| Gender | 0.26<br>(0.20) |
| Sample | 0.07<br>(0.30) |
| BDI (prop. score) x Gender | −0.28<br>(0.58) |
| BDI (prop. score) x Sample | −0.15<br>(0.92) |
| Gender x Sample | −0.35<br>(0.50) |
| BDI (prop. score) x Gender x Sample | 1.27<br>(2.13) |
| Num.Obs. | 294 |
| R<sup>2</sup> | 0.016 |
| R<sup>2</sup> Adj. | −0.008 |
| AIC | 694.5 |
| BIC | 727.6 |
| Log.Lik. | −338.246 |
| F | 0.655 |
| RMSE | 0.76 |

+ $p < 0.1$ , \* $p < 0.05$ , \*\* $p < 0.01$ , \*\*\* $p < 0.001$

**Table 11:** Multi-level model analysis coefficients and standard errors for effort in reward-seeking. BAI (prop. score) = anxiety score on the Beck Anxiety Inventory. Proportion scores are scores divided by total possible score. AIC = Akaike information criterion, BIC = Bayesian information criterion, ICC = intraclass correlation, RMSE = root mean squared error.

|  | Multi-level model: BAI pred. Effort |
| --- | --- |
| (Intercept) | 5.27***<br>(0.26) |
| BAI (prop. score) | 1.44*<br>(0.71) |
| Block | −0.87***<br>(0.02) |
| Gender | 1.30***<br>(0.31) |
| Sample | 1.08*<br>(0.49) |
| BAI (prop. score) x Block | 0.00<br>(0.05) |
| BAI (prop. score) x Gender | −3.35***<br>(0.93) |
| Block x Gender | −0.03<br>(0.02) |
| BAI (prop. score) x Sample | −2.92+<br>(1.52) |
| Block x Sample | −0.12**<br>(0.04) |
| Gender x Sample | −2.05**<br>(0.76) |
| BAI (prop. score) x Block x Gender | 0.15*<br>(0.07) |
| BAI (prop. score) x Block x Sample | 0.39**<br>(0.13) |
| BAI (prop. score) x Gender x Sample | 3.62<br>(4.81) |
| Block x Gender x Sample | 0.22***<br>(0.06) |
| BAI (prop. score) x Block x Gender x Sample | −0.79+<br>(0.42) |
| SD (Intercept participant) | 1.30 |
| SD (Observations) | 1.98 |
| Num.Obs. | 37677 |
| R2 Marg. | 0.282 |
| R2 Cond. | 0.498 |
| AIC | 159770.0 |
| BIC | 159923.7 |
| ICC | 0.3 |
| RMSE | 1.98 |

**Table 12:** Multi-level model analysis coefficients and standard errors for effort in reward-seeking. BDI (prop. score) = depression score on the Beck Depression Inventory II. Proportion scores are scores divided by total possible score. AIC = Akaike information criterion, BIC = Bayesian information criterion, ICC = intraclass correlation, RMSE = root mean squared error.

|  | Multi-level model: BDI pred. Effort |
| --- | --- |
| (Intercept) | 5.64***<br>(0.29) |
| BDI (prop. score) | 0.15<br>(0.82) |
| Block | −0.88***<br>(0.02) |
| Gender | 0.73*<br>(0.35) |
| Sample | 0.44<br>(0.53) |
| BDI (prop. score) x Block | 0.03<br>(0.06) |
| BDI (prop. score) x Gender | −1.01<br>(1.03) |
| Block x Gender | 0.01<br>(0.03) |
| BDI (prop. score) x Sample | −0.51<br>(1.64) |
| Block x Sample | −0.07<br>(0.04) |
| Gender x Sample | −1.29<br>(0.89) |
| BDI (prop. score) x Block x Gender | 0.00<br>(0.08) |
| BDI (prop. score) x Block x Sample | 0.17<br>(0.13) |
| BDI (prop. score) x Gender x Sample | 1.45<br>(3.81) |
| Block x Gender x Sample | 0.14+<br>(0.07) |
| BDI (prop. score) x Block x Gender x Sample | −0.25<br>(0.32) |
| SD (Intercept participant) | 1.32 |
| SD (Observations) | 1.98 |
| Num.Obs. | 37677 |
| R2 Marg. | 0.276 |
| R2 Cond. | 0.499 |
| AIC | 159796.9 |
| BIC | 159950.6 |
| ICC | 0.3 |
| RMSE | 1.98 |

**Table 13:** Linear model analysis coefficients and standard errors for breakpoint in reward-seeking. BAI (prop. score) = anxiety score on the Beck Anxiety Inventory. Proportion scores are scores divided by total possible score. AIC = Akaike information criterion, BIC = Bayesian information criterion, Log. Lik. = log likelihood, RMSE = root mean squared error.

|  | Linear model: BAI pred. Sensitivity (d') |
| --- | --- |
| (Intercept) | 113.69***<br>(3.21) |
| BAI (prop. score) | 7.21<br>(8.83) |
| Gender | 6.73+<br>(3.84) |
| Sample | −2.79<br>(6.05) |
| BAI (prop. score) x Gender | −13.91<br>(11.57) |
| BAI (prop. score) x Sample | 9.61<br>(18.66) |
| Gender x Sample | 3.79<br>(9.34) |
| BAI (prop. score) x Gender x Sample | −3.59<br>(59.57) |
| Num.Obs. | 294 |
| R2 | 0.019 |
| R2 Adj. | −0.005 |
| AIC | 2507.8 |
| BIC | 2540.9 |
| Log.Lik. | −1244.881 |
| F | 0.786 |
| RMSE | 16.70 |

+ p < 0.1, \* p < 0.05, \*\* p < 0.01, \*\*\* p < 0.001

**Table 14:**
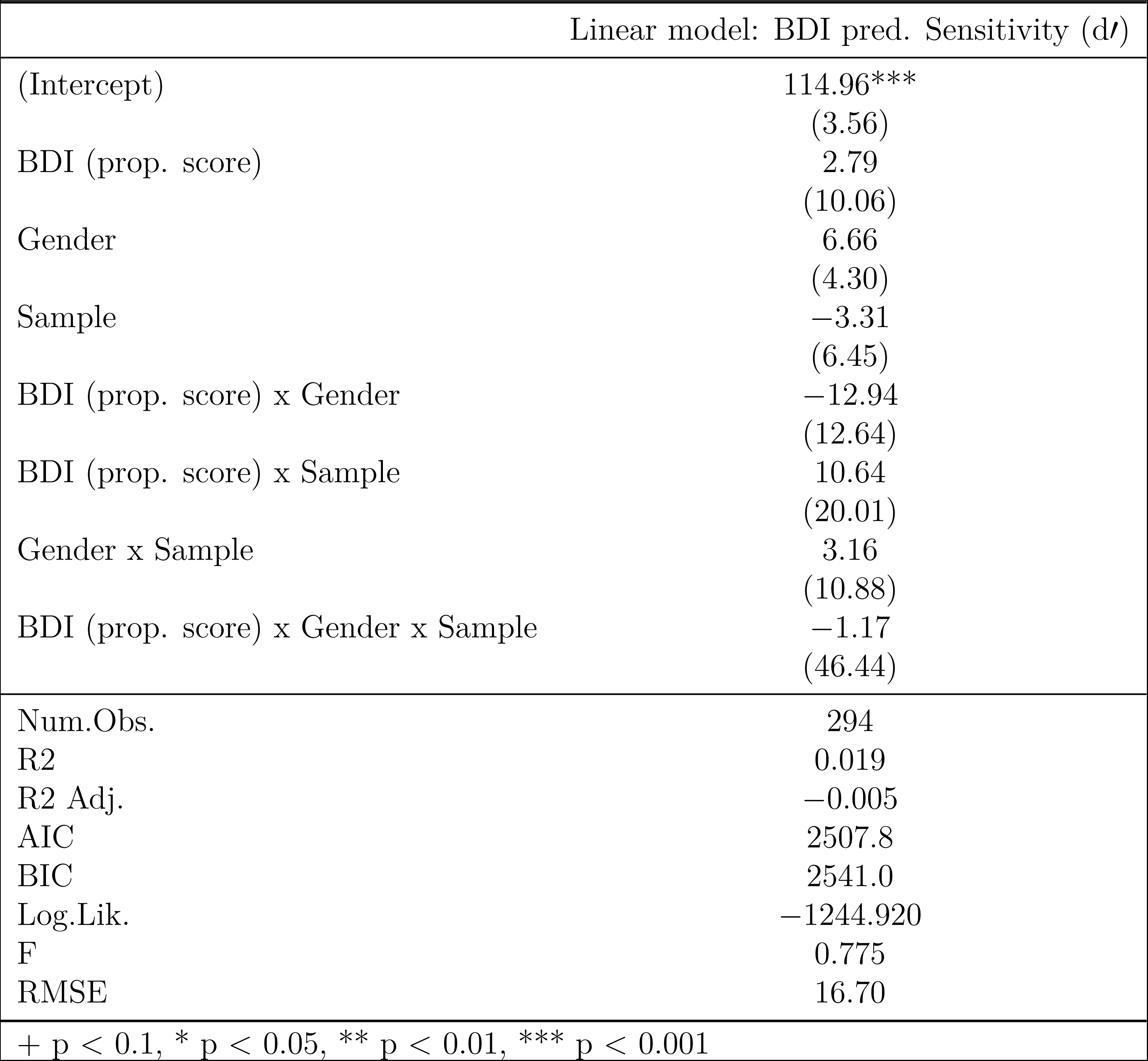
Linear model analysis coefficients and standard errors for breakpoint in reward-seeking. BDI (prop. score) = depression score on the Beck Depression Inventory II. Proportion scores are scores divided by total possible score. AIC = Akaike information BIC = Bayesian information criterion, Log. Lik. = log likelihood, RMSE = root ared error.

|  | Linear model: BDI pred. Sensitivity (d') |
| --- | --- |
| (Intercept) | 114.96***<br>(3.56) |
| BDI (prop. score) | 2.79<br>(10.06) |
| Gender | 6.66<br>(4.30) |
| Sample | −3.31<br>(6.45) |
| BDI (prop. score) x Gender | −12.94<br>(12.64) |
| BDI (prop. score) x Sample | 10.64<br>(20.01) |
| Gender x Sample | 3.16<br>(10.88) |
| BDI (prop. score) x Gender x Sample | −1.17<br>(46.44) |
| Num.Obs. | 294 |
| R2 | 0.019 |
| R2 Adj. | −0.005 |
| AIC | 2507.8 |
| BIC | 2541.0 |
| Log.Lik. | −1244.920 |
| F | 0.775 |
| RMSE | 16.70 |

+ p < 0.1, \* p < 0.05, \*\* p < 0.01, \*\*\* p < 0.001

