## Supplementary material for "Gender impacts the relationship between mood disorder symptoms and effortful avoidance performance"

### Avoidance task

We first compared participants' accuracy on active and inhibitory avoidance trials by gender using a 2x2 within-between ANOVA (Trial Type x Gender) (Table 1). A main effect revealed that, on average, participants were more accurate on inhibitory than on active trials ($F(1, 521) = 405.38$, p $<$ .001, $\eta^2 = 0.25$; Table 2). There was a main effect of gender ($F(1, 521) = 5.79$, $p = .02$, $\eta^2 = < .01$) as well as a gender interaction such that men had higher accuracy than women on active but not inhibitory avoidance trials ($F(1, 521) = 4.98$, $p = .03$, $\eta^2 < .01$).

Depression and anxiety symptoms were not significantly associated with inhibitory or active avoidance accuracy (Table 2). However, given observed gender differences in both depression/anxiety disorder symptoms and task performance, we conducted a series of moderation analyses using the PROCESS macro (Hayes, 2017) as implemented in `bruceR` (Bao, 2021) to examine whether gender moderated the relationship between accuracy and anxiety and depressive symptoms (Fig. 1B). We found that gender moderated the relationship between anxiety symptoms (as measured in BAI scores) and inhibitory avoidance accuracy ($F(1, 540) = 4.24$, $p = .04$; Fig. 1A). Increased BAI scores were associated with lower inhibitory avoidance accuracy in females but not males; this relationship was not observed for BDI scores. For the full results of the moderation analysis, see Table 3.

**Table 1**
*ANOVA of accuracy on active and inhibitory avoidance trials. BAI = Beck Anxiety Inventory, BDI = Beck Depression Inventory.*

| Effect | $df_n$ | $df_d$ | $F$ | $p$ | sig. | $\eta^2$ |
|---|---|---|---|---|---|---|
| Gender | 1 | 521 | 5.79 | 0.016 | * | $< .01$ |
| Trial Type | 1 | 521 | 405.38 | $< .001$ | *** | 0.25 |
| Gender:Trial Type | 1 | 521 | 4.98 | 0.026 | * | $< .01$ |

**Table 2**
*Relationship between accuracy and mood disorder symptoms on active and inhibitory avoidance trials. BAI = Beck Anxiety Inventory, BDI = Beck Depression Inventory.*

| Trial Type | Comparison | df | $t$ | $p$ | sig. |
|---|---|---|---|---|---|
| Active avoidance | BDI~accuracy | 540 | -0.32 | 0.75 | |
| | BAI~accuracy | 540 | -1.36 | 0.17 | |
| Inhibitory avoidance | BDI~accuracy | 540 | -0.87 | 0.38 | |
| | BAI~accuracy | 540 | -0.73 | 0.47 | |

**Table 3**
*Gender moderation of relationship between accuracy and mood disorder symptoms on active and inhibitory avoidance trials. BAI = Beck Anxiety Inventory, BDI = Beck Depression Inventory.*

| Trial Type | Comparison | $df_n$ | $df_d$ | $t$ | $p$ | sig. |
|---|---|---|---|---|---|---|
| Active avoidance | Gender moderation of BDI~accuracy | 1 | 540 | 0.01 | 0.92 | |
| | Gender moderation of BAI~accuracy | 1 | 540 | 0.29 | 0.59 | |
| Inhibitory avoidance | Gender moderation of BDI~accuracy | 1 | 540 | 1.03 | 0.31 | |
| | Gender moderation of BAI~accuracy | 1 | 540 | 4.24 | 0.04 | * |

To further examine the relationship between accuracy and anxiety and depressive symptoms, we tested whether BAI scores predicted accuracy in men and women separately. Outside of the moderation analysis after Bonferroni correction, BAI scores overall predicted inhibitory but not active avoidance performance accuracy (Fig. 1B, ($t(542) =$ -2.93, $p = .06$, $r =$ -0.12)). Additionally, higher BAI scores predicted lower inhibitory avoidance performance accuracy in women but not men (Fig. 1B; ($t(259) =$ -3.71, $p = .01$, $r =$ -0.22, *95% CI* [-0.35, -0.09])).

We also conducted an interactive regression analysis to investigate whether the relationship between anxiety symptoms and active and inhibitory avoidance accuracy differed as a function of the level of depressive symptoms (Fig. 3). In active avoidance, the relationship between anxiety symptoms (BAI scores) and accuracy for all participants was a function of their level of depressive symptoms (BDI scores) ($t(540) =$ -2.64, $p = .01$); however, this was not the case in inhibitory avoidance ($t(540) =$ -0.88, $p = .38$). As such, only those with higher levels of depressive symptoms were impaired in active avoidance, and not those with higher levels of anxiety symptoms but lower levels of depressive symptoms.

We first compared participants' accuracy on active and inhibitory reward-seeking trials by gender (Fig. 2A) using a 2x2 within-between ANOVA (Trial Type x Gender) (Table 4). A main effect revealed that, on average, participants were more accurate on inhibitory than on active trials ($F(1, 292) = 58.86$, p $< .001$, $\eta^2 = 0.07$). There was no main effect of gender ($F(1, 292) = 2.61$, $p = .11$, $\eta^2 = 0.01$) but there was a gender interaction such that men had higher accuracy than women on active but not inhibitory reward-seeking trials ($F(1, 292) = 7.61$, $p = .01$, $\eta^2 = 0.01$).

Given observed gender differences in both depressive/anxiety symptoms and task performance, we again conducted a series of moderation analyses to see whether gender moderated the relationship between accuracy and anxiety and depressive symptoms. Anxiety was associated with reduced active reward seeking accuracy in the total sample. However, we found no gender moderation of the relationship between anxiety or depressive symptoms and active or inhibitory reward-seeking accuracy, and BDI scores did not interact with active or inhibitory reward-seeking accuracy (Table 5, Table 6, Fig. 2A).

To further elucidate the relationship between accuracy and anxiety and depressive symptoms, we evaluated whether BAI and BDI scores predicted accuracy in men and women separately. After Bonferroni correction, BAI and BDI scores did not predict overall active or inhibitory reward-seeking performance in either men or women (Fig. 2B). We also conducted an interactive regression analysis to investigate whether the relationship between anxiety symptoms and active and inhibitory reward-seeking accuracy differed as a function of their level of depressive symptoms (Fig. 4). In active reward-seeking, the relationship between anxiety symptoms (BAI scores) and accuracy for all participants was not a function of their level of depressive symptoms (BDI scores) ($t(306) = 0.45$, $p = .66$); this was also not the case in inhibitory reward-seeking ($t(306) = -0.78$, $p = .44$).

**Table 4**
*ANOVA of accuracy on active and inhibitory reward-seeking trials.*

| Effect | $df_n$ | $df_d$ | $F$ | $p$ | sig. | $\eta^2$ |
|---|---|---|---|---|---|---|
| Gender | 1 | 292 | 2.61 | 0.107 | | 0.01 |
| Trial Type | 1 | 292 | 58.86 | < .001 | *** | 0.07 |
| Gender:Trial Type | 1 | 292 | 7.61 | 0.006 | ** | 0.01 |

**Table 5**
*Relationship between accuracy and mood disorder symptoms on active and inhibitory reward-seeking trials. BAI = Beck Anxiety Inventory, BDI = Beck Depression Inventory II.*

| Trial Type | Comparison | df | $t$ | $p$ | sig. |
|---|---|---|---|---|---|
| Active reward-seeking | BDI~accuracy | 306 | -0.52 | 0.61 | |
| | BAI~accuracy | 306 | -2.56 | 0.01 | * |
| Inhibitory reward-seeking | BDI~accuracy | 306 | -0.54 | 0.59 | |

| Trial Type | Comparison | df | *t* | *p* | sig. |
|---|---|---|---|---|---|
| | BAI~accuracy | 306 | -0.83 | 0.41 | |

**Table 6**
*Gender moderation of relationship between accuracy and mood disorder symptoms on active and inhibitory reward-seeking trials. BAI = Beck Anxiety Inventory, BDI = Beck Depression Inventory II.*

| Trial Type | Comparison | $df_n$ | $df_d$ | *t* | *p* | sig. |
|---|---|---|---|---|---|---|
| Active reward-seeking | Gender moderation of BDI~accuracy | 1 | 306 | 0.56 | 0.46 | |
| | Gender moderation of BAI~accuracy | 1 | 306 | 3.30 | 0.07 | |
| Inhibitory reward-seeking | Gender moderation of BDI~accuracy | 1 | 306 | 0.12 | 0.73 | |
| | Gender moderation of BAI~accuracy | 1 | 306 | 0.00 | 0.96 | |

Bao H-W-S (2021) bruceR: Broadly useful convenient and efficient r functions.

Hayes AF (2017) Introduction to mediation, moderation, and conditional process analysis: A regression-based approach. Guilford publications.

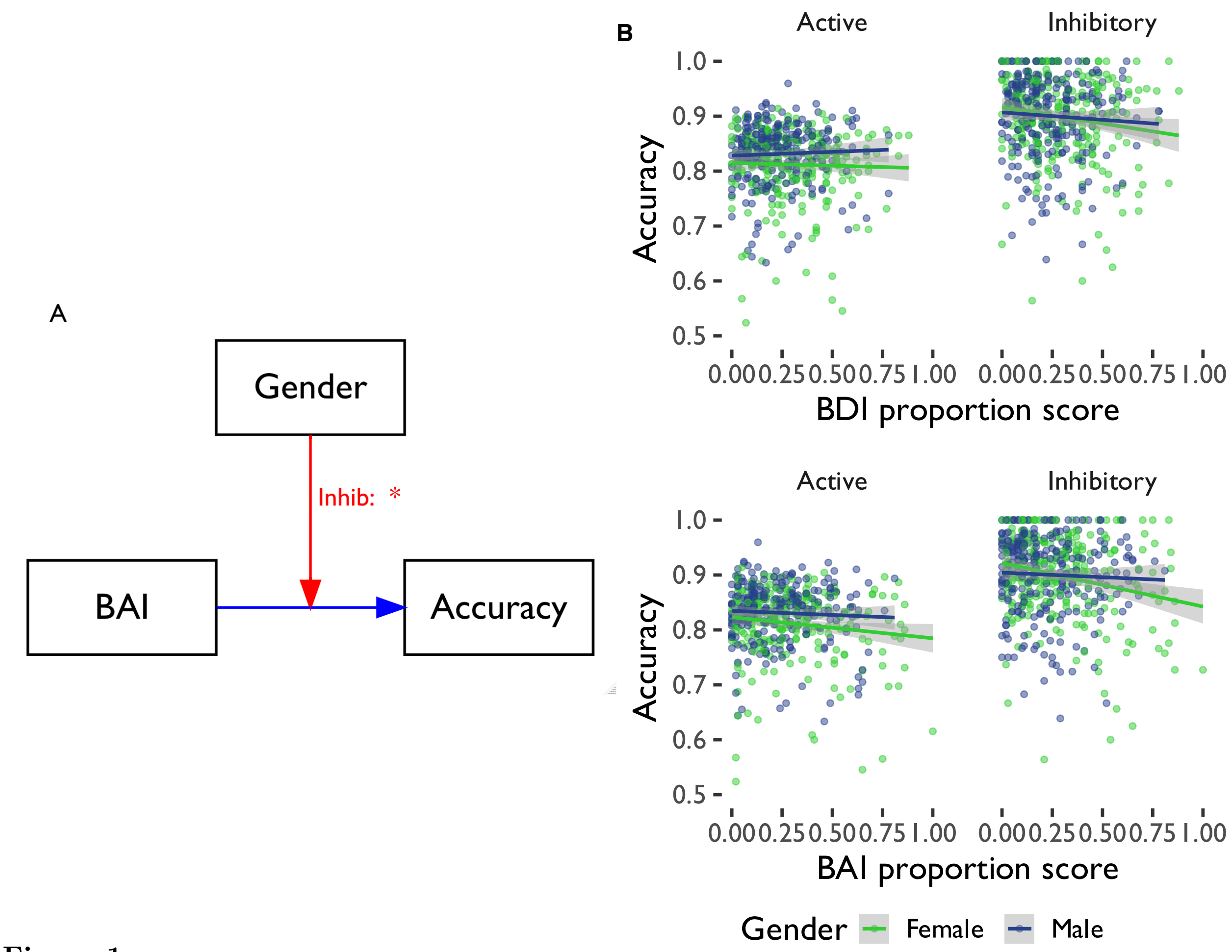


**Figure 1**

*(A) Moderation of the relationship between accuracy and anxiety (BAI) proportion scores by gender in active and inhibitory avoidance. Gender significantly moderated the relationship between anxiety symptoms (BAI proportion scores) and inhibitory avoidance accuracy. (B) Accuracy by gender on active and inhibitory avoidance. BAI = Beck Anxiety Inventory. Proportion scores are scores divided by total possible score.*

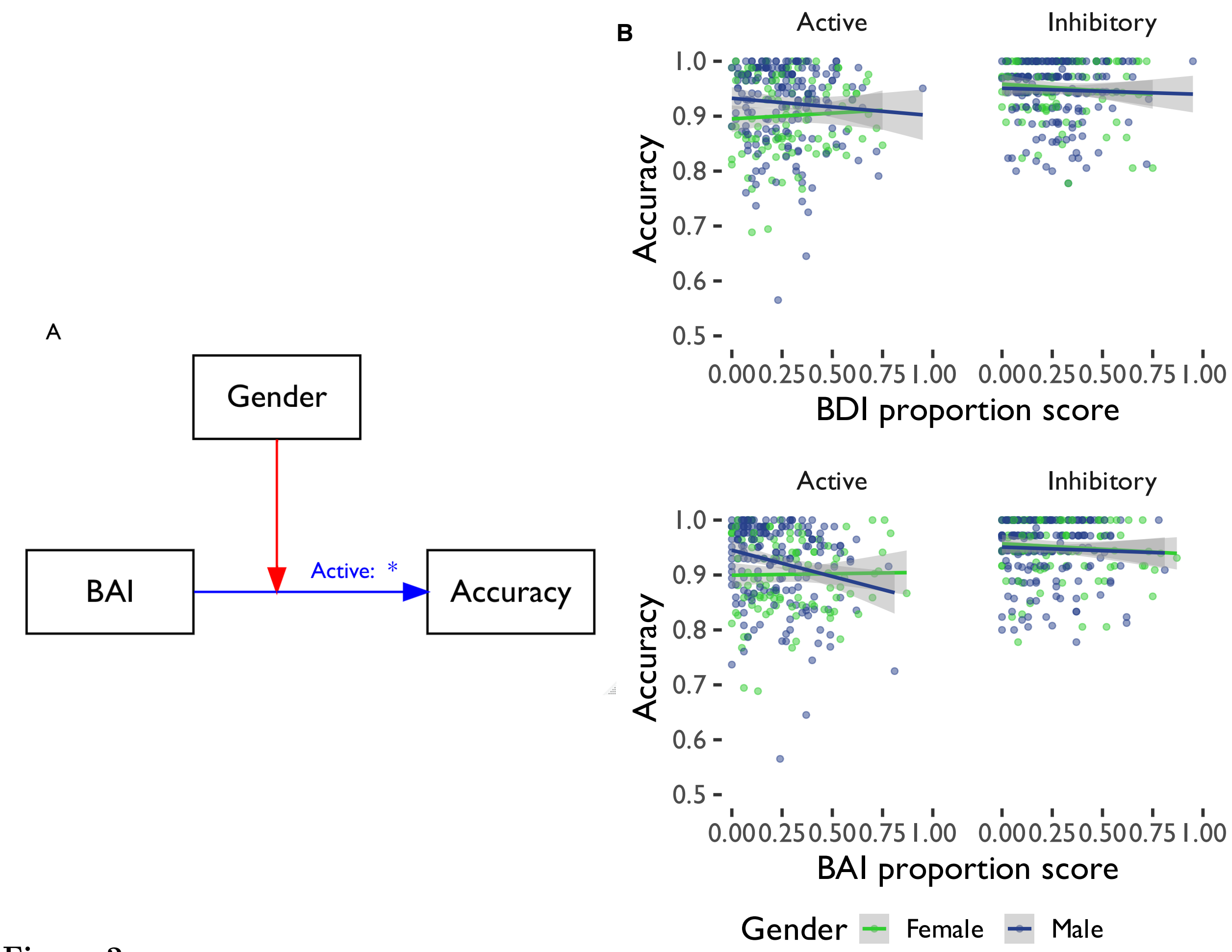


**Figure 2**

*(A) Moderation of the relationship between accuracy and anxiety (BAI) proportion scores by gender on active and inhibitory reward-seeking. Anxiety symptoms (BAI proportion scores) were significantly associated with active reward-seeking accuracy. (B) Accuracy by gender on active and inhibitory reward-seeking. BAI = Beck Anxiety Inventory. Proportion scores are scores divided by total possible score.*

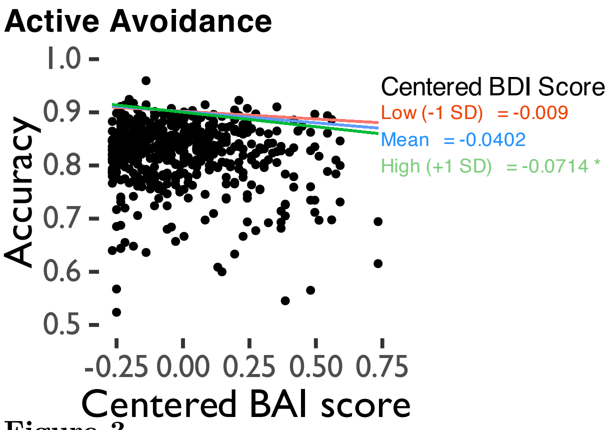


**Figure 3**

*Interactive regression analysis output for active and inhibitory avoidance. BAI = Beck Anxiety Inventory, BDI = Beck Depression Inventory II. Numbers on right indicate coefficients for regression lines at each level of BDI score (centred 1 standard deviation below the mean, at the mean, and 1 standard deviation above the mean).*

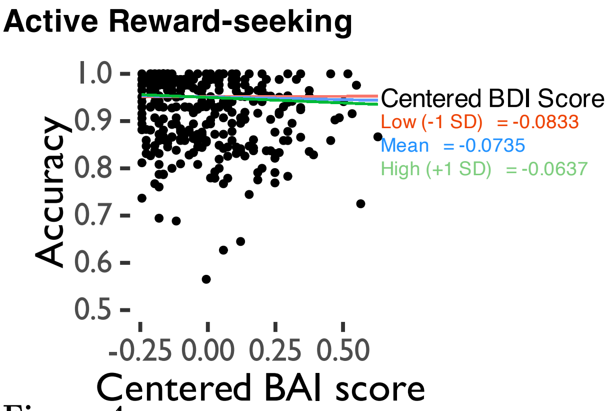


**Figure 4**

*Interactive regression analysis output for active and inhibitory reward-seeking. BAI = Beck Anxiety Inventory, BDI = Beck Depression Inventory II. Numbers on right indicate coefficients for regression lines at each level of BDI score (centred 1 standard deviation below the mean, at the mean, and 1 standard deviation above the mean).*
